# CRISPR activation screens identify core protein-dependent regulation of heparan sulfate sulfation and ligand specificity

**DOI:** 10.64898/2026.06.29.735380

**Authors:** Jack C. Moore, Haruki Takeuchi, Caitlien Nguyen, Chin Huang, Digantkumar Chapla, Amrita Basu, Zhangjie Wang, Jian Liu, Kelley W. Moremen, Ryan J. Weiss

**Affiliations:** Complex Carbohydrate Research Center, University of Georgia, Athens, GA; Department of Biochemistry and Molecular Biology, University of Georgia, Athens, GA; Glycan Therapeutics Corp, Raleigh, NC; Division of Chemical Biology and Medicinal Chemistry, Eshelman School of Pharmacy, University of North Carolina, Chapel Hill, NC

**Keywords:** heparan sulfate, proteoglycan, syndecan, glycosaminoglycan, CRISPR activation, antithrombin

## Abstract

Heparan sulfate proteoglycans (HSPGs) are essential cell surface and extracellular matrix glycoconjugates that mediate diverse biological processes through interactions between their heparan sulfate (HS) chains and extracellular ligands. While HS sulfation patterning is known to dictate ligand specificity, how cells control HS assembly to regulate these interactions remains incompletely understood. To systematically identify genetic modifiers of HS-protein interactions, we performed genome-wide CRISPR activation (CRISPRa) screens in HEK293T cells using binding of antithrombin (AT), which selectively recognizes 3-*O*-sulfated HS motifs, or the *N*-sulfation-specific antibody 10E4 as functional readouts. Strikingly, the screens revealed proteoglycan core proteins as key modulators of HS function. In particular, syndecan-1 (SDC1) emerged as a preferential enhancer of AT binding compared to other syndecan family members. Targeted upregulation of syndecan family members increased total HS levels, but only SDC1 enhanced AT binding. Structural and enzymatic analyses demonstrated that SDC1-associated HS chains contain elevated 6-*O*-sulfation and serve as superior substrates for 3-*O*-sulfotransferases relative to SDC2-associated HS chains. Additionally, SDC1 exhibited slower cell surface recovery, which was blocked by cycloheximide treatment, consistent with extended trafficking and biosynthetic processing. Overall, these findings indicate that proteoglycan core protein identity influences HS sulfation patterning and ligand-binding specificity and trafficking kinetics may contribute to core protein-dependent regulation of HS modification.

## Introduction

Heparan sulfate proteoglycans (HSPGs) are key cell surface and extracellular matrix (ECM) components that regulate morphogen gradients, growth factor signaling, cell adhesion, and tissue organization. HSPGs are comprised of a core protein covalently decorated with one or more heparan sulfate (HS) chains at specific serine residues within consensus linkage regions [1, 2]. The function of HSPGs is primarily determined by the structure and sulfation pattern of their HS chains, which create diverse binding sites for extracellular ligands [3]. Certain interactions are highly selective; for example, the serine protease inhibitor antithrombin (AT) binds to rare 3-*O*-sulfated pentasaccharide motifs within HS or the anticoagulant drug, heparin, which is a highly sulfated form of HS (Fig. 1A) [4, 5]. Binding leads to an AT conformational change and potent inhibition of thrombin and factor Xa in the coagulation cascade [6]. Other ligands, such as members of the fibroblast growth factor (FGF) family, vascular endothelial growth factors (VEGFs), and their receptors, engage HS more flexibly to modulate cell signaling, thus relying on sulfation density and domain organization rather than strict sequence requirements [3, 7, 8]. Thus, HSPG function depends not only on HS abundance but on precise sulfation patterning that governs ligand selectivity and downstream signaling. Despite extensive characterization of HS-binding ligands and biosynthetic enzymes, the mechanisms by which cells regulate the assembly of ligand-permissive HS motifs remain poorly understood.

**Figure 1.**
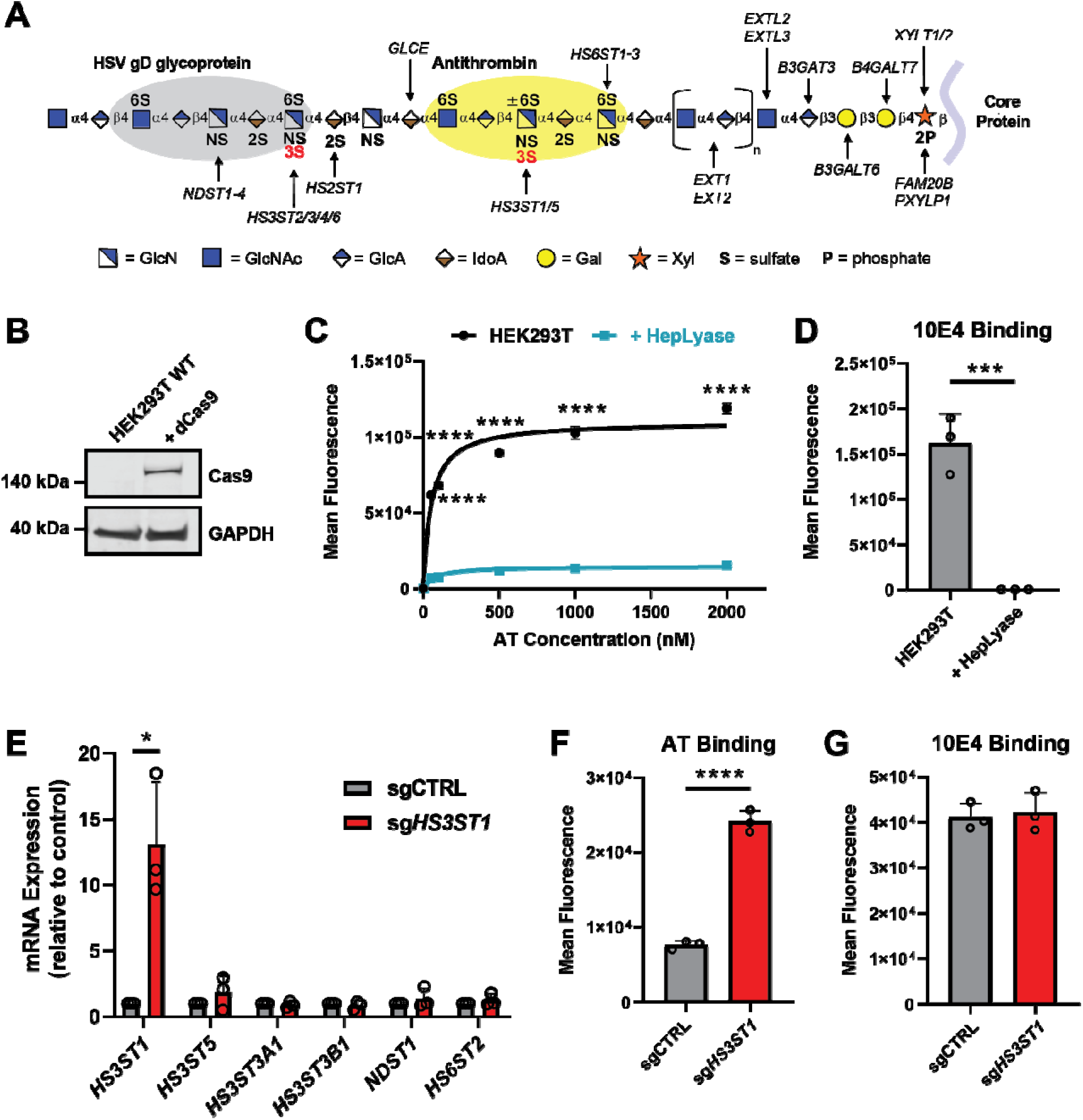
CRISPR activation selectively enhances heparan sulfate sulfation and ligand binding in HEK293T cells. **(A)** Schematic representation of heparan sulfate (HS) biosynthesis. HS is elongated and sulfated while attached via a tetrasaccharide linker to a proteoglycan core protein. Seven different 3-*O* sulfotransferases (HS3ST1-6) can add sulfate groups to carbon-3 of *N*-acetylglucosamine (GlcNAc) and glucosamine (GlcN) residues. **(B)** Western blot showing ectopic dCas9-VP64 expression in transduced HEK293T cells. GAPDH was used as a loading control. **(C)** Flow cytometry analysis of antithrombin (AT) binding to HEK293T cells with and without pre-treatment with heparan lyase I-III (HepLyase) over a gradient of AT concentrations. **(D)** Flow cytometry analysis of 10E4 binding of HEK293T cells pre-treated with and without heparan lyases (I-III). **(E)** qPCR analysis of mRNA expression of HS3ST1 and related HS biosynthetic genes in sg*HS3ST1* activation cell line compared to non-targeting control sgRNA (sgCTRL) cells. Flow cytometry analysis of **(F)** AT and **(G)** 10E4 binding for sg*HS3ST1* cells compared to sgCTRL cells. Data are presented as mean ± SD (n = 3 independent experiments), \*\*\**p*<0.0001, \*\*\**p*<0.001, \*\**p*<0.01, \**p*<0.05 by two-sided t-test.

Following proper folding in the endoplasmic reticulum, proteoglycan core proteins transit to the Golgi apparatus, where HS chains are assembled at specific serine residues and sequentially modified by glycosyltransferases, epimerases, and sulfotransferases. These coordinated reactions generate distinct *N*-acetylated (NA) and *N*-sulfated (NS) domains decorated with 2-*O*-, 6-*O*-, and rare 3-*O*-sulfation, producing structurally heterogeneous HS chains capable of diverse ligand interactions (Fig. 1A) [1, 9]. While the HS enzymatic machinery is well characterized, the mechanisms that govern formation of ligand-selective motifs, the arrangement of domains, and their context on specific proteoglycan core proteins remain largely unexplored. In particular, it is unclear whether proteoglycan core proteins are passive scaffolds for HS attachment or whether they can influence the structure and functional output of their attached chains. Emerging evidence suggests that factors such as Golgi trafficking dynamics [10–13], enzyme distribution and dynamics [14–16], and substrate availability [5, 17, 18] may also influence HS domain formation, raising the possibility that core protein identity contributes to sulfation patterning. Understanding these mechanisms is critical for connecting HS composition to selective ligand interactions in development, homeostasis, and disease.

Advances in CRISPR screening technologies over the past decade have provided expanded strategies for genome-wide dissection of regulatory mechanisms for biological processes [19, 20]. Our laboratory and others have utilized genome-wide CRISPR screening approaches to identify novel regulatory factors controlling mammalian glycosylation, including both *N*-glycans [21–23] and GAGs [24–26]. To systematically identify regulators of HS-ligand interactions, we reasoned that a gain-of-function approach would be particularly informative since many HS biosynthetic enzymes are expressed at low levels, exhibit cell type-restricted expression, or functionally redundant, limiting the sensitivity of conventional loss-of-function screens. In this study, we therefore employed genome-wide CRISPRa screening coupled to functional ligand-binding readouts and flow cytometry to identify genetic determinants that enhance selective HS-protein interactions. For screening, we utilized antithrombin (AT) binding at the cell surface to detect 3-*O*-sulfated HS motifs and the *N*-sulfation-specific antibody 10E4 as a complementary measure of overall HS levels, respectively. Unexpectedly, this approach identified the HSPG syndecan-1 (SDC1) as a preferential enhancer of AT binding relative to other syndecan family members. Comparative analyses of HS derived from SDC1 and SDC2 demonstrate core protein-dependent differences in sulfation patterning, enzymatic susceptibility to 3-*O* sulfation, and biosynthetic kinetics. Together, our results reveal that core protein identity is associated with differences in HS assembly, shaping selective ligand interactions at the cell surface.

## Results

### Genome-wide CRISPR activation screens identify regulators of HS-ligand interactions

To identify genetic factors that regulate functional HS-protein interactions, we generated HEK293T cells stably expressing a catalytically dead Cas9 (dCas9) fused to a VP64 transcriptional activation domain and transduced them with the Calabrese genome-wide sgRNA library [27] (Fig. 1B). HEK293T cells were selected for our studies since they grow rapidly in culture, are amendable to CRISPR engineering, present modestly sulfated cell surface HS [28–30], and express many of the known HS biosynthetic enzymes and proteoglycan core proteins (Suppl. Fig. 1A). Prior to screening, we validated HS-dependent cell surface binding of AT, which selectively recognizes 3-*O*-sulfated HS pentasaccharide motifs [31, 32], and 10E4, an HS-specific antibody targeting *N*-sulfated hybrid domains [33]. Flow cytometry demonstrated that AT and 10E4 binding was abolished by heparan lyase treatment, confirming specificity of both probes for cell surface HS chains (Fig. 1C-D). To confirm proper CRISPRa functionality in HEK293T-dCas9 cells, we stably expressed a single guide RNA (sgRNA) targeting *HS3ST1* (sgHS3ST1), which encodes for the primary 3-*O* sulfotransferase responsible for generating the AT binding site within heparin/HS chains [34]. These cells exhibited robust and specific transcriptional upregulation of *HS3ST1*, as confirmed by quantitative PCR, with minimal expression changes of a subset of related HS biosynthetic genes (Fig. 1E). Functionally, upregulation of *HS3ST1* increased cell surface binding of AT by ∼3-fold compared to wildtype control cells without altering 10E4 binding (Fig. 1F-G).

We next performed genome-wide CRISPRa screening by transducing HEK293T-dCas9 cells at a low multiplicity of infection (MOI ∼0.3) with the Calabrese sgRNA library followed by fluorescence-activated cell sorting (FACS) based on increased binding of either AT or 10E4 using fluorescence-activated cell sorting (FACS) (Fig. 2A). Cells within the top 1% of AT or 10E4 binding were isolated from triplicate biological replicates, and sgRNA representation was quantified by next-generation sequencing and analysis using PinAPL-Py [35]. Analysis of recovered sgRNA sequences from the AT and 10E4 binding screens showed strong enrichment of a small subset sgRNAs with highly skewed averaged read count distributions and Gini coefficients of ∼0.97 and ∼0.96, respectively, across the high AT and 10E4 binding samples (Suppl. Fig. 1B-C). In the 10E4 screen, the top hit was syndecan 2 (*SDC2*), a known cell surface heparan sulfate proteoglycan (HSPG) [36], thus validating the screening approach as a readout of increased HS abundance (Fig. 2B). Notably, multiple heparan sulfate-related genes were enriched in this population, including additional proteoglycans (*SDC1, SDC4, GPC1*) and HS-modifying enzymes, including *HS3ST4*. Gene ontology analysis identified *HS-GAG biosynthesis* as the top enriched pathway, as well as additional pathways related to hedgehog signaling, protein-protein interactions, and hemopoiesis (Fig. 2D). Notably, canonical HS polymerases such as EXT1/EXT2 were not enriched, indicating that HS chain elongation is not rate-limiting under these conditions [1] and/or promoter-dependent activation constraints. For the AT-binding screen, *HS3ST1* was the top hit, consistent with its established role in generating AT-binding motifs within heparin/HS chains (Fig. 2C). Additional enriched genes included the HSPGs, syndecan-1 (*SDC1*) and glypican-1 (*GPC1*), *HS3ST4*, and *PAPSS1,* which biosynthesizes the sulfate donor PAPS needed for HS modification [25]. Pathway analysis of the top hits from the AT high-binding screen again revealed *HS-GAG biosynthesis* as the top enriched pathway, including other pathways related to hedgehog signaling and neuroligand interactions (Fig. 2E). Together, these results validate the 10E4 and AT screening approaches and identify new potential pathways and genetic factors involved in regulating heparan sulfate presentation and function.

**Figure 2.**
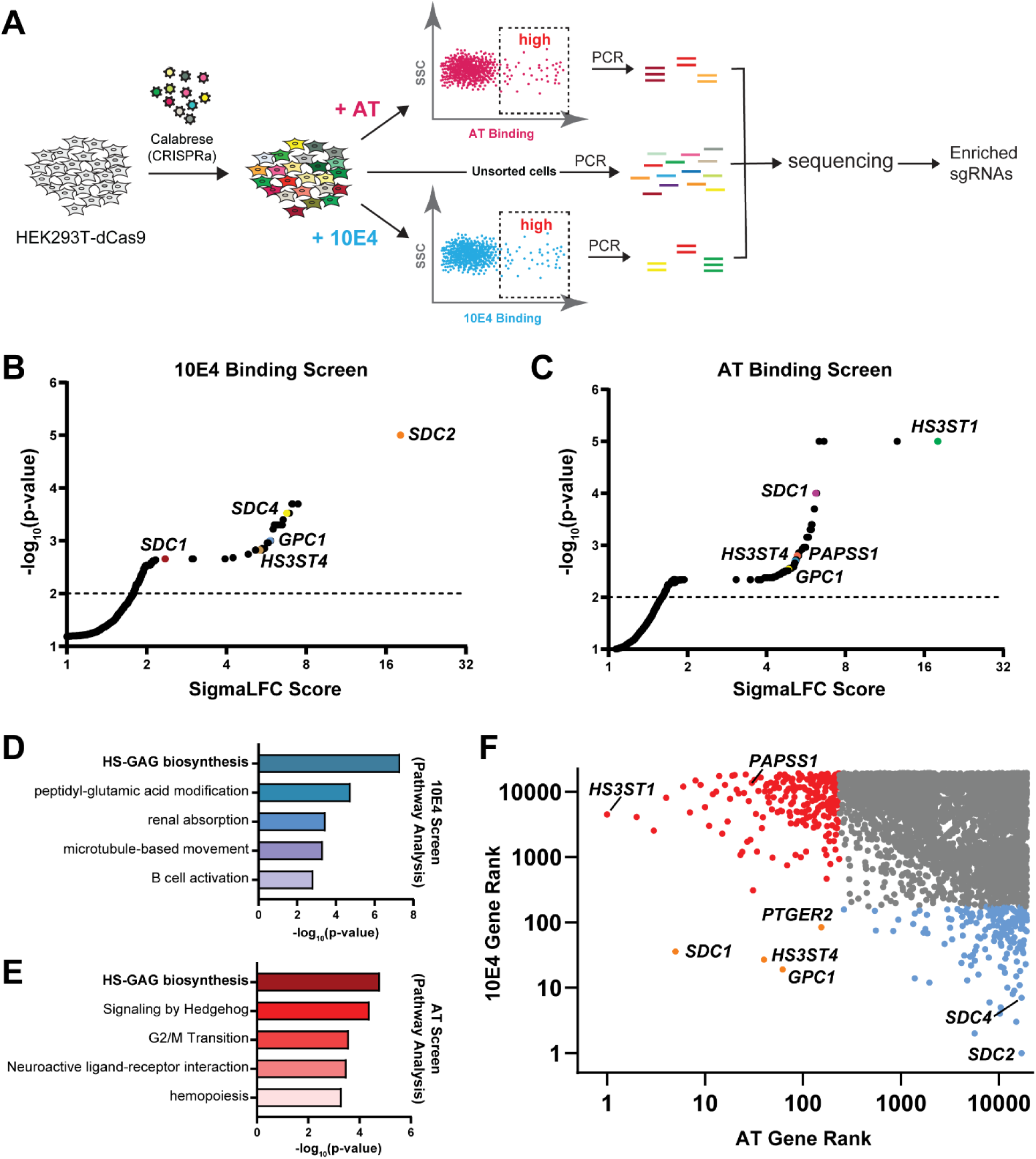
CRISPR activation screens identify syndecan proteoglycans with distinct and shared AT and 10E4 binding profiles. **(A)** HEK293T-dCas9 cells were transduced with the genome-wide Calabrese sgRNA library and screened for high AT (top) or high 10E4 (bottom) binding (n = 3 biological replicates). Unsorted cells were collected to assess sgRNA library coverage. **(B)** Scatterplot of the averaged gene enrichment results from the high 10E4 binding screens (n = 3 biological replicates). Gene hits were ranked based on significance (p-value) and enrichment (SigmaLFC) compared to unsorted control cell population. Red and blue data points represent genes significantly enriched in the AT and 10E4 sorted populations, respectively. Selected genes involved in HSPG biosynthesis are annotated on the graph. **(C)** Scatterplot of the averaged gene enrichment results from the high AT binding screens (n = 3 biological replicates). **(D-E)** Gene ontology analysis of the top significantly enriched genes (p ≤ 0.05) in 10E4 or AT sorted populations, respectively. **(F)** Comparison of ranked enriched genes from 10E4 sorted and AT sorted populations. Genes rankings were assigned based on SigmaLFC scores. Red and blue data points mark genes significantly enriched in the AT and 10E4 sorted populations, respectively (p ≤ 0.01). Orange dots highlight significantly enriched genes across both screening approaches. Select genes involved in HSPG biosynthesis are highlighted on the graph.

Due to the shared specificity of the 10E4 and AT probes for HS chains, we next compared the results from both screens, which revealed strong shared enrichment (*p* ≤ 0.01) of four genes: *SDC1*, *HS3ST4*, *GPC1,* and *PTGER2* (Fig. 2F). These hits included two HSPGs (*SDC1*, *GPC1*), a HS 3-O-sulfotransferase (*HS3ST4*), and a prostaglandin E2 receptor (*PTGER2*) that has not been previously implicated in HSPG biosynthesis. A striking divergence emerged within the syndecan family, with *SDC1* enriched in both screens and *SDC2* and *SDC4* selectively enriched only in the 10E4 screen. These screening results suggested that while increased expression of select proteoglycans can increase overall cell surface HS abundance, only a subset may promote the formation of HS species capable of binding AT.

### Activation of syndecan family proteoglycans reveals differential HS functional outputs

To mechanistically investigate how proteoglycan core protein identity influences HS structure and/or ligand binding, we focused on the syndecan family, which are type I single-pass transmembrane proteoglycans that are present on most mammalian cell surfaces. Also known as “hybrid” proteoglycans, this family of proteoglycans have been shown to carry HS and/or chondroitin/dermatan sulfate (CS/DS) glycosaminoglycan (GAG) chains at distinct serine residues within their extracellular intrinsically disordered ectodomain [37] (Fig. 3A). Structurally, the syndecans share similarities in the topology of their transmembrane domain and cytoplasmic tail; however, their ectodomains differ significantly in size and amino acid sequence [38]. To establish a controlled system for comparing syndecan-dependent HS outputs, we first generated individual CRISPR activation lines using specific sgRNAs for all four syndecans (sgSDC1, sgSDC2, sgSDC3, sgSDC4) in HEK293T-dCas9 cells. Initial attempts to utilize the VP64 CRISPRa system gave variable and weak activation for single sgRNA targets, thus the SAM CRISPRa system [39] was used for downstream individual syndecan activation experiments. Upon stable cell line generation, we confirmed strong and selective upregulation of each respective syndecan transcript, with minimal cross-regulation of the other family members (Fig. 3B). To confirm that changes in mRNA expression resulted in increased levels of proteoglycan protein presentation at the cell surface, we quantified cell surface levels for each syndecan using specific commercial antibodies and flow cytometry. We detected a significant increase in each syndecan at the cell surface in each respective activation cell line compared to non-targeting control cells (Fig. 3C-F). Comparatively, we found the greatest fold-increase (∼15-fold) in cell surface SDC2 protein levels, with an average of ∼10-fold across the syndecan family, compared to wild-type control cells.

**Figure 3.**
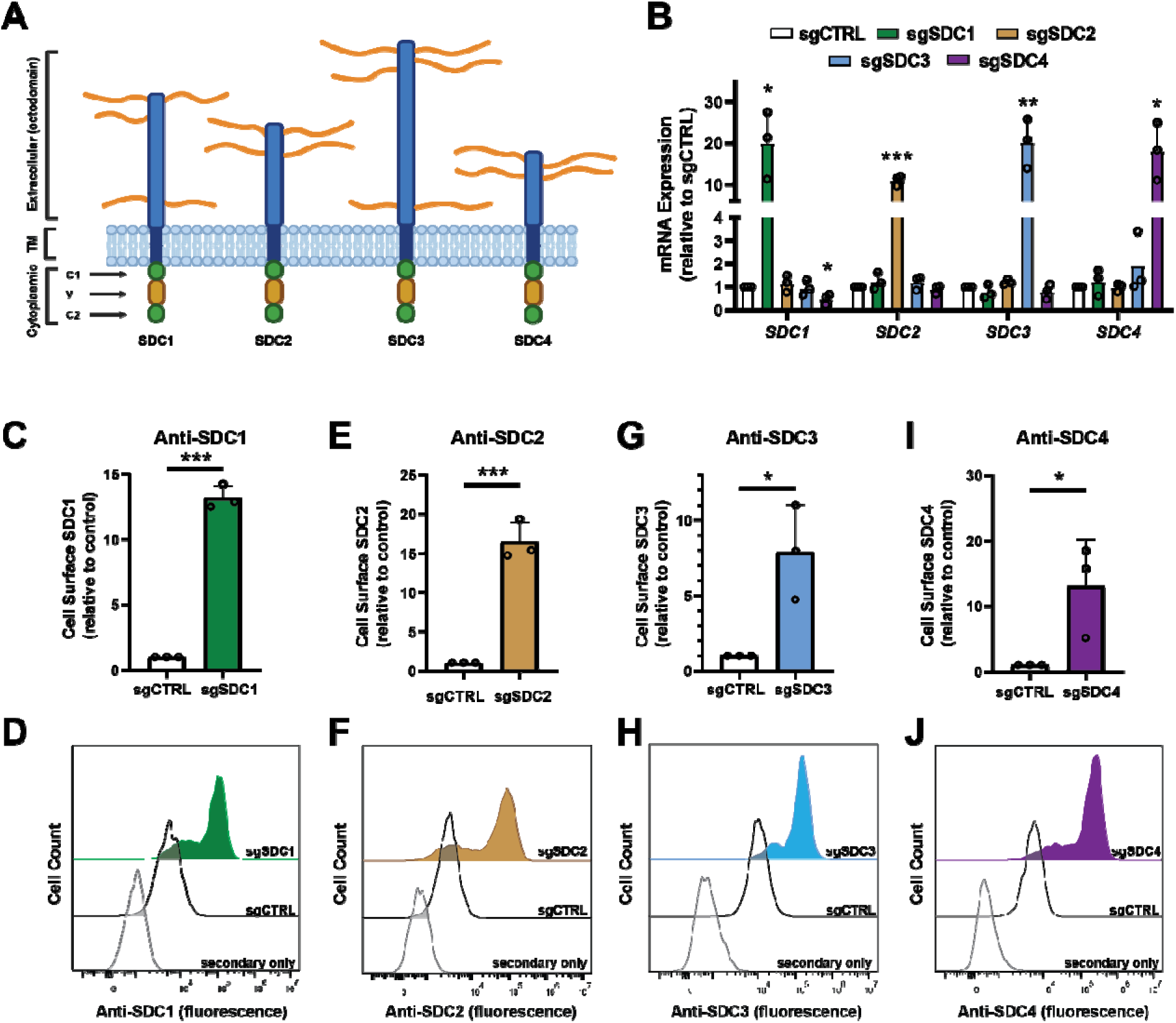
CRISPR-mediated activation of syndecan expression selectively enhances proteoglycan presentation at the cell surface. **(A)** Cartoon depicting shared topology of the four syndecan proteoglycan family members, which are single-pass transmembrane proteins containing an extracellular, transmembrane, and cytoplasmic domain, respectively. Their cytoplasmic domain consists of 2 conserved regions (C1, C2) and a variable region (V), while their extracellular ectodomains differ in length and are diversely modified with HS and CS/DS polysaccharide chains (orange lines) at specific sites. **(B)** mRNA expression levels of *SDC1-4* in HEK293T CRISPRa cell lines. Quantitative bar graphs and histogram overlays of flow cytometry analysis of **(C-D)** SDC1, **(E-F)** SDC2, **(G-H)** SDC3, and **(I-J)** SDC4 protein levels at the cell surface with specific antibodies in HEK293T CRISPRa cell lines. Data are presented as mean ± SD (n = 3 independent experiments), \*\*\**p*<0.001, \*\**p*<0.01, \**p*<0.05 by two-sided t-test. Panel **(A)** was created in BioRender. Moore, J.C. https://BioRender.com/jr7ozgd (2026).

We next profiled GAG presentation and ligand binding across the syndecan activation lines. All syndecan activation cell lines showed increased total cell surface HS levels, measured by 10E4 binding, with SDC2 exhibiting the largest increase (∼4-fold), consistent with its identification as the top hit in the 10E4 screen (Fig. 4A). SDC1 also significantly increased HS levels (∼3-fold), while SDC3 and SDC4 showed more modest (∼1.5-fold) effects. We further assessed HSPG presentation and HS attachment sites using the antibody 3G10, which detects a neoepitope generated upon pre-treatment with a cocktail of heparin lyase enzymes [33]. All four syndecan activation lines showed a significant increase in 3G10 binding, with SDC2 and SDC4 displaying the largest increase (Fig. 4B). Importantly, despite broadly increased cell surface HS abundance, only SDC1 activation resulted in a significant increase in AT binding, whereas SDC2 and SDC4 showed no measurable change and SDC4 exhibited a slight decrease in AT binding (Fig. 4C). This dissociation confirmed that increased HS levels alone were insufficient to generate AT-binding motifs and that core protein identity influences HS sulfation patterning. In addition, we assessed cell surface CS levels using the anti-chondroitin sulfate antibody, CS-56 [40]. This analysis revealed a modest (∼2-fold) increase in CS-56 binding upon *SDC1-3* activation, with SDC4 showing a ∼10-fold increase in CS presentation, suggesting that SDC4 is primarily modified by CS chains at the cell surface in the HEK293T cells (Fig. 4D). Collectively, these results demonstrated that syndecans differentially regulate HS and CS presentation at the cell surface. Notably, SDC1 and SDC2 exhibited opposing phenotypes, where SDC2 upregulation gave a robust increase in total HS abundance without enhancing AT binding, and SDC1 selectively promoted the formation of AT binding. Based on this functional divergence, we focused subsequent analyses on directly comparing SDC1 and SDC2.

**Figure 4.**
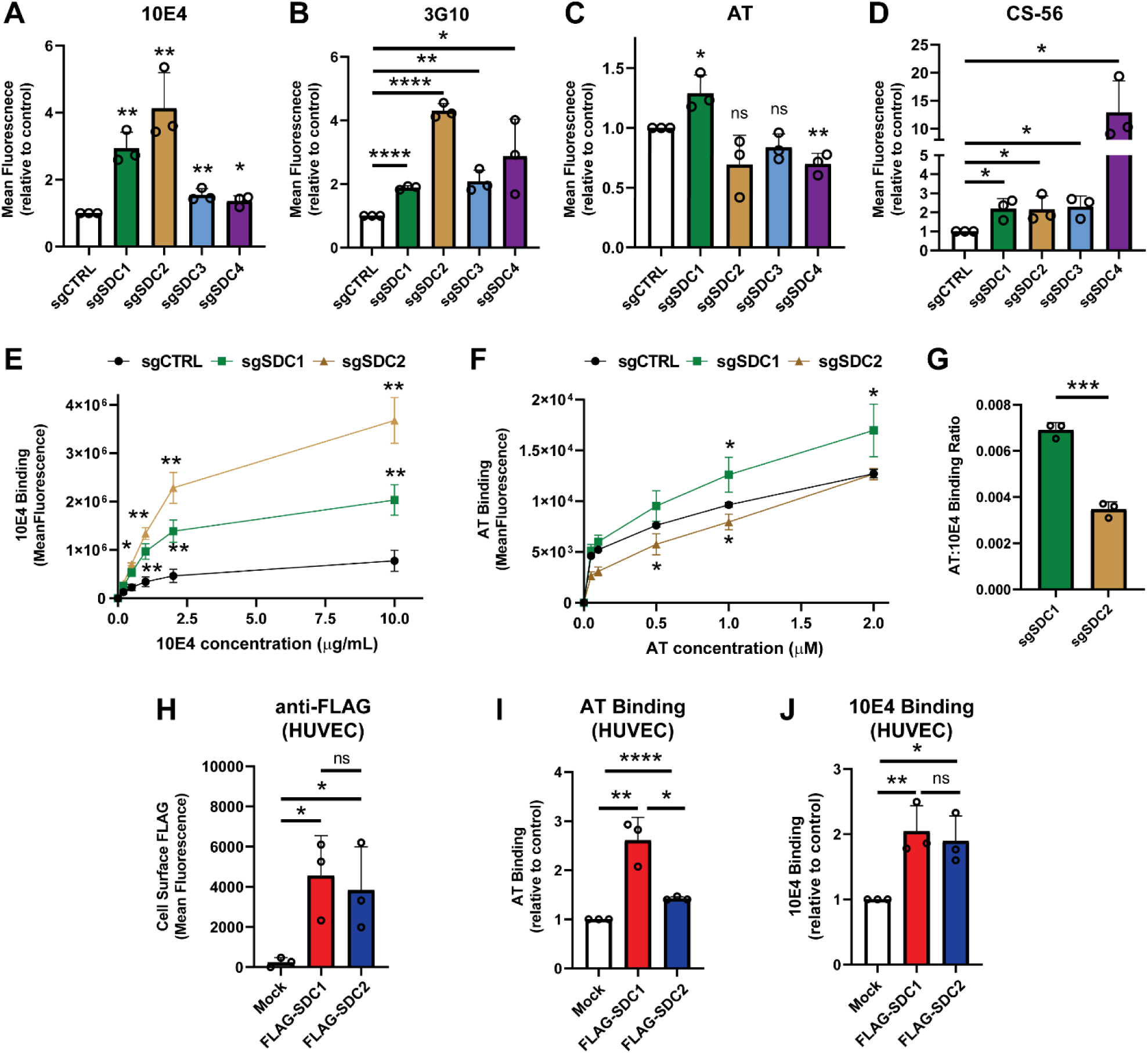
Upregulation of *SDC1* selectively enhances AT binding in HEK293T cells and primary human endothelial cells. **(A)** Flow cytometry analysis of 10E4 binding in CRISPRa cell lines versus non-targeting control cells. **(B)** Flow cytometry analysis of 3G10 binding in CRISPRa cell lines pre-treated with heparan lyases I-III. **(C)** Flow cytometry analysis of AT binding in CRISPRa cell lines versus non-targeting control cells. **(D)** Flow cytometry analysis of CS-56 binding in CRISPRa cell lines versus non-targeting control cells. **(E)** 10E4 and **(F)** AT binding across a concentration gradient in sgSDC1 and sgSDC2 activation cell lines versus non-targeting control (sgCTRL) cells. **(G)** Quantification of the ratio of AT to 10E4 binding at 2 µM and 10 µg/mL concentrations, respectively. **(H)** Quantification of cell surface N-terminal FLAG-tag levels by flow cytometry for primary human endothelial cells (HUVECs) transfected with N-FLAG-SDC1 or N-FLAG-SDC2 constructs. **(I)** Flow cytometry analysis of cell surface AT binding in transfected HUVECs. **(J)** Flow cytometry analysis of cell surface 10E4 binding in transfected HUVEC cells. Data are presented as mean ± SD (n = 3 independent experiments), \*\*\*\**p*<0.0001, \*\*\**p*<0.001, \*\**p*<0.01, \**p*<0.05 by two-sided t-test.

### SDC1 and SDC2 differentially regulate HS ligand binding specificity

To directly compare HS functional outputs, we analyzed ligand binding in SDC1 and SDC2 activation cells across a range of ligand concentrations. SDC2 activation consistently resulted in a pronounced increase in 10E4 binding with no corresponding increase in AT interaction over a concentration range (Fig. 4E-F). In contrast, SDC1 activation produced a more modest increase in total HS with a marked increase in AT binding over a range of concentrations (Fig. 4E-F). This difference in cell surface binding was highlighted further by plotting the ratio of AT to 10E4 signal at their highest respective concentrations, which revealed a significantly higher AT to 10E4 ratio in sgSDC1 cells relative to sgSDC2 (Fig. 4G). The binding behavior of additional HS binding proteins was quantified by flow cytometry, which revealed an increase in FGF1 and VEGF_165_ binding similar to 10E4, while FGF2 was unchanged in both sgSDC1 and sgSDC2 cell lines, suggesting that the HS structural differences driven by SDC1 versus SDC2 do not uniformly enhance all HS-ligand interactions (Suppl. Fig. 2A-C). Additional flow cytometry experiments confirmed no compensation for SDC1 or SDC2 presentation at the cell surface in both sgSDC1 and sgSDC2 lines (Suppl. Fig. 2D-E).

To validate these findings in a more physiologically relevant model, we transfected N-terminal FLAG-tagged SDC1 and SDC2 constructs into primary human umbilical vein endothelial cells (HUVECs) and assessed AT and 10E4 binding versus mock transfected control cells. Importantly, HUVECs are known to express *HS3ST1* and produce anticoagulant HS which is present at the endothelial surface [41, 42], making them a biologically meaningful system to evaluate the consequence of syndecan-associated HS patterning in the presence of the canonical AT-binding site. We first validated that both constructs were expressed at comparable levels on the cell surface, as measured with an anti-FLAG antibody and flow cytometry (Fig. 4H). Consistent with our HEK293T results, SDC1 overexpression produced a significant increase in AT binding (∼3-fold) compared to SDC2 (Fig. 4I) while both constructs provided a similar increase in 10E4 binding (Fig. 4J), thus showing that differential promotion of AT-binding HS by SDC1 is observed in a physiologically relevant cell context.

### SDC1-associated HS exhibits enhanced sulfation and increased susceptibility to 3-*O* modification

To determine the molecular basis for the observed differences in HS-protein interactions, we next analyzed the composition of GAG chains associated with SDC1 and SDC2, respectively, via expression of recombinant ectodomains of SDC1 and SDC2 (eSDC1, eSDC2). The recombinant proteins were produced using an HEK293 expression system [43] and analyzed for GAG compositional analysis (Fig. 5A). Total GAGs were isolated from eSDC1 and eSDC2 and subjected to disaccharide and 3-*O*-sulfated tetrasaccharide analyses using LC-MS methods, as previously described [44, 45]. HS disaccharide analysis revealed that eSDC1-derived HS chains contained significantly higher levels of specific 2-*O*- and 6-*O*-sulfated disaccharides relative to eSDC2, including increases in ΔUA2S-GlcNAc, ΔUA-GlcNAc6S, ΔUA-GlcNS6S, and ΔUA2S-GlcNS6S species (Fig. 5B-C, Suppl. Tables 1-2). Analysis of 3-*O* sulfated tetrasaccharides indicated elevated levels of ΔUA-GlcNAc6S-GlcA-GlcNS3S6S (D0N6-G0S9) and ΔUA-GlcNS6S-GlcA-GlcNS3S6S (D0S6-G0S9) species in eSDC1 HS, which are both enriched within the AT pentasaccharide motif of pharmaceutical heparins [44] but are at relatively lower abundance in HEK293-derived syndecan HS chains (Fig. 5D, Suppl. Table 3).

**Figure 5.**
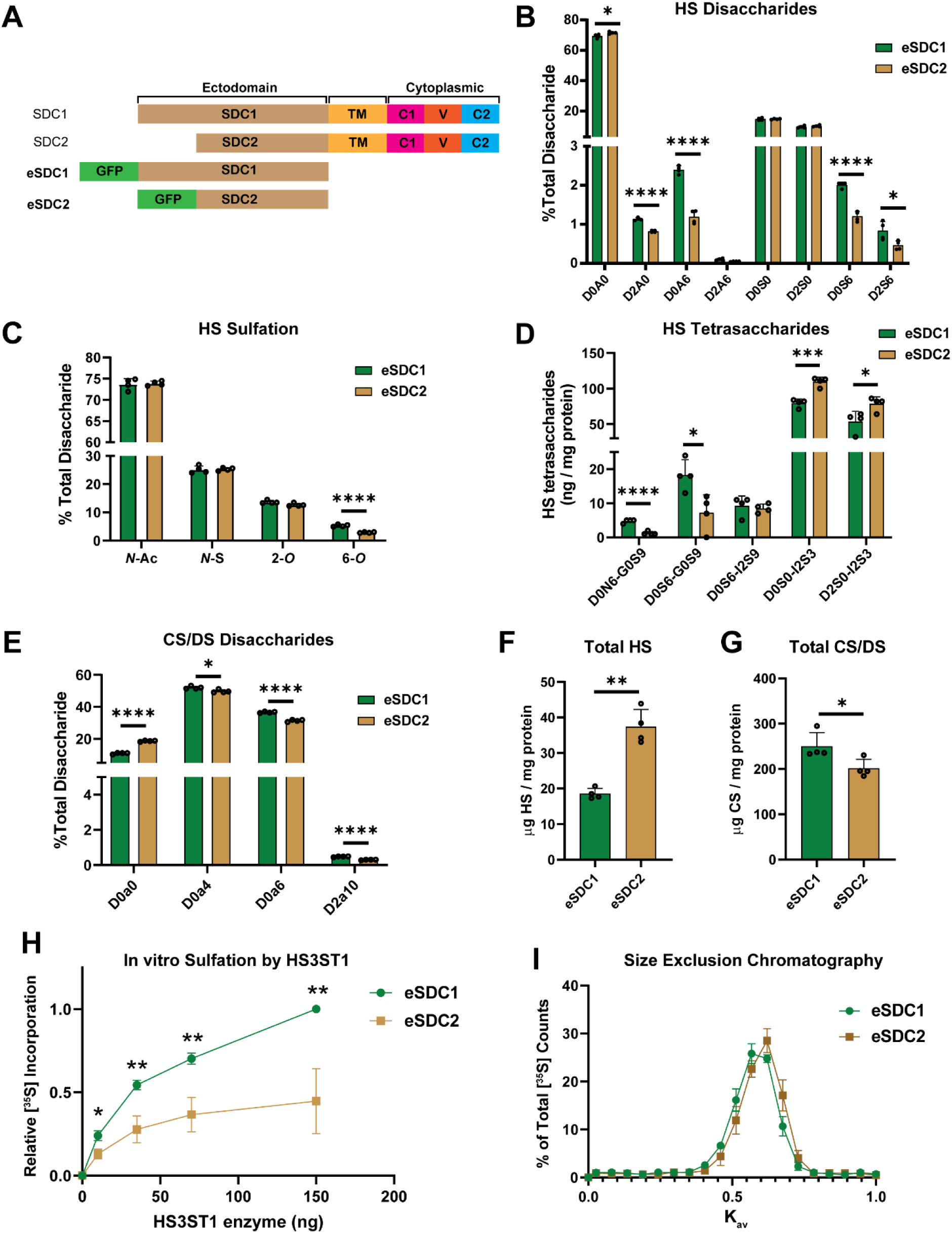
SDC1-associated HS exhibits enhanced sulfation and increased susceptibility to 3-*O* sulfation. **(A)** Schematic of full length SDC1 and SDC2 compared to N-terminal GFP-tagged syndecan ectodomain constructs (eSDC1, eSDC2). **(B)** LC-MS disaccharide analysis of HS chains isolated from eSDC1 and eSDC2. **(C)** Comparison of total abundance of sulfation residues in isolated HS. The disaccharide structure code is described in Suppl. Table 1 and [85]. **(D)** LC-MS 3-*O*-sulfated tetrasaccharide analysis of HS chains isolated from eSDC2 and eSDC2. **(E)** LC-MS disaccharide analysis of CS/DS chains isolated from eSDC1 and eSDC2. **(F)** In vitro 3-*O* sulfation assay with recombinant HS3ST1 and HS chains isolated from eSDC1 and eSDC2. Counts were normalized to the maximum signal measured within a given experiment. **(G)** LC-MS quantification of total HS isolated from eSDC1 and eSDC2. **(H)** LC-MS quantification of total CS/DS isolated from eSDC1 and eSDC2. **(I)** Size exclusion chromatography of HS chains [^35^S]-labeled by recombinant HS3ST1 using a Sepharose 6B column. Data are presented as mean ± SD (n = 3-4 independent experiments), \*\*\*\**p*<0.0001, \*\*\**p*<0.001, \*\**p*<0.01, \**p*<0.05 by two-sided t-test.

Notably, the most abundant tetrasaccharide species, D0S0-I2S3, was slightly increased in eSDC2 HS, highlighting that the abundance of AT- and gD-type (associated with herpes simplex virus recognition [46]) 3-*O*-sulfated subdomains are distinct between syndecan-derived HS chains. Analysis of CS/DS chains isolated from eSDC1 and eSDC2 revealed elevated 4-*O* and 6-*O* sulfated disaccharides (D0a4, D0a6, D2a10) for eSDC1 (Fig 5E, Suppl. Table 4). Both syndecan ectodomains contained more CS/DS species relative to HS (Fig. 5F-G), with eSDC2 showing higher HS content, consistent with elevated 3G10 and 10E4 levels found in sgSDC2 cells above (Fig. 4A-B). Together, these results demonstrated enhanced total GAG sulfation in eSDC1-derived GAG chains, with distinct differences in overall GAG content.

Since HS is synthesized and sequentially modified within the Golgi apparatus, with 3-*O* sulfation representing one of the final steps in this pathway, we next asked whether HS derived from eSDC1 and eSDC2 differed in their capacity to serve as substrates for 3-*O* sulfotransferases. To test this directly, we adapted *in vitro* 3-*O* sulfation assays using [^35^S]-PAPS and recombinant HS3ST1, as previously described [47, 48]. Early attempts to utilize intact syndecan ectodomains were unsuccessful, thus we purified GAGs via beta-elimination and anion exchange chromatography along with removal of CS/DS chains using chondroitinase ABC. Across a gradient of enzyme concentrations, HS purified from eSDC1 was consistently modified to a greater extent (∼3-fold) than HS from eSDC2, indicating that eSDC1-associated HS chains contain a higher abundance of precursor sequences permissive for 3-*O* sulfation by HS3ST1 (Fig. 5H). This trend was reproduced using recombinant HS3ST5 (Suppl. Fig. 2F). To determine whether differences in HS chain length could account for altered enzyme activity, HS3ST1 modified [^35^S]HS chains were analyzed by size exclusion chromatography. HS derived from both eSDC1 and eSDC2 exhibited nearly identical elution profiles, with an average K_av_ of ∼0.59, indicating comparable chain lengths (Fig. 5I). Collectively, our findings demonstrate differential incorporation of 3-*O* sulfate groups due to intrinsic fine structural differences in the HS chains associated with SDC1 versus SDC2 ectodomains.

### SDC1 and SDC2 exhibit distinct biosynthetic trafficking kinetics

Given the increased overall sulfation observed in SDC1-associated GAG chains, we hypothesized that SDC1 and SDC2 may differ in their intracellular trafficking kinetics, thereby altering the time available for HS chain modification during Golgi transit. Directly measuring Golgi residence and trafficking rates for individual endogenous proteoglycans is technically challenging. We therefore developed a cell surface recovery assay following proteolytic removal of proteoglycans as a functional readout to measure their combined biosynthetic and trafficking kinetics. To establish baseline surface levels of SDC1 and SDC2, HEK293T cells were gently lifted using EDTA and analyzed by flow cytometry. To remove surface-exposed ectodomains, parallel cultures were treated with trypsin, immediately neutralized, and either collected on ice as a trypsin-treated baseline or replated to allow surface recovery (Fig. 6A). Reappearance of syndecans at the cell surface, which reflected de novo synthesis, Golgi processing, and trafficking through the secretory pathway, was monitored at 2-, 4-, 6-, and 8-hour time points to plot initial rates of recovery. Due to the observed long cell surface recovery times of these proteoglycans [49], we focused on obtaining initial recovery rates in live cells. Strikingly, SDC2 reappeared at the plasma membrane ∼2-fold faster than SDC1 (Fig. 6B-C). Although SDC2 recovered to ∼65% of baseline surface levels within 8 hours, SDC1 recovered to only approximately ∼40% of its baseline level over the same period. Importantly, pre- and concurrent treatment of cells with cycloheximide to inhibit protein synthesis abolished cell surface recovery of both proteoglycans (Fig. 6B), demonstrating that surface repopulation following trypsin treatment was dependent on new protein synthesis rather than recycling of endocytosed proteoglycans or mobilization of pre-formed intracellular pools.

**Figure 6.**
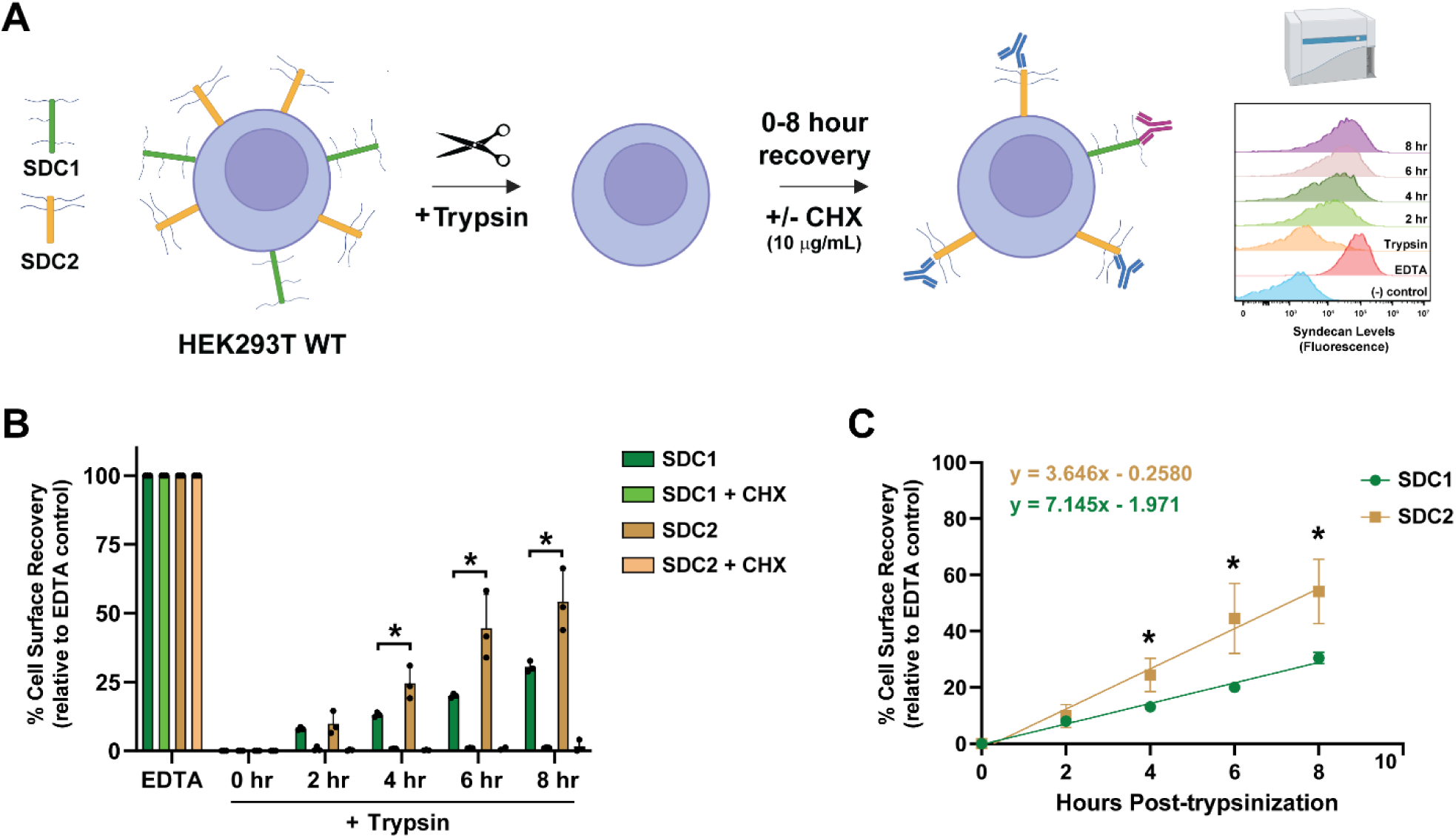
SDC1 exhibits reduced biosynthetic trafficking kinetics. **(A)** Schematic of the cell surface recovery assays. Cells were either gently detached with PBS (+ 10 mM EDTA) to establish baseline cell surface syndecan levels or were treated with trypsin to proteolytically remove surface-exposed ectodomains, then were replated and monitored for proteoglycan surface recovery at 2-8 hours by flow cytometry. **(B)** SDC1 and SDC2 cell surface recovery assays in HEK293T cells. Cells were treated with trypsin and the recovery of either SDC1 or SDC2 to the cell surface in the presence or absence of cycloheximide (CHX) was monitored via flow cytometry. Recovery levels were normalized to cells treated with EDTA as a baseline. Cycloheximide-treated cells gave a significant reduction in signal (*p*<0.0001). **(C)** Recovery of proteoglycans to the cell surface were plotted to determine linear fit and slope of the recovery rate over time. Data are presented as mean ± SD (n = 3 independent experiments), \**p*<0.05 by two-sided t-test. Panel **(A)** was created in BioRender. Moore, J.C. https://BioRender.com/zzvh1hc (2026).

## Discussion

Heparan sulfate biosynthesis has long been described as a non-template-driven process governed principally by the expression, activity, and localization of Golgi-resident glycosyltransferases and sulfotransferases, with the proteoglycan core protein serving largely as a passive scaffold for chain attachment. While prior work has suggested that core protein sequence context may influence GAG chain type specification and/or polymerization [50–53], the extent to which core protein identity shapes sulfation patterning and ligand-binding specificity has remained an open question. In this study, we utilized genome-wide CRISPRa screening with functional ligand-binding readouts, which unexpectedly revealed proteoglycan core protein identity as a modulator of HS sulfation. Specifically, CRISPRa screens identified syndecan-1 (SDC1) as a selective enhancer of AT binding, while other syndecan family members primarily increased total HS abundance. Biochemical dissection of SDC1- and SDC2-associated HS demonstrated that SDC1 carries chains enriched in 6-*O* sulfation and altered 3-*O-*sulfated moieties, that SDC1-derived HS is a superior substrate for 3-*O* sulfotransferases, and that SDC1 exhibits slower cell surface recovery kinetics consistent with extended biosynthetic processing. Together, these findings support a model in which the proteoglycan core protein contributes to shaping the HS modification landscape, with implications for how cells generate and regulate ligand-selective HS sequences.

CRISPRa screening provided a robust gain-of-function discovery platform to interrogate HS biosynthetic regulation. Specifically, transcriptional upregulation of endogenous proteoglycan genes enabled the detection of proteoglycan-specific regulation of HS-protein interactions at the cell surface. The complementary use of two functionally distinct probes, AT and 10E4, revealed an unexpected dichotomy within the syndecan proteoglycan subfamily, with SDC2 as the top hit exclusively in the 10E4 screen, while SDC1 was the top overlapping hit across both screens. This divergence indicated that increased HS abundance and increased HS functional complexity are separable properties, and that core protein identity is a key variable controlling HS-ligand interactions. Notably, the canonical HS polymerases EXT1 and EXT2 or HS initiating enzyme EXTL3 were not enriched in either screen (Fig. 2), suggesting that HS biosynthetic output is likely constrained at the level of substrate availability and/or proteoglycan core protein expression. This aligns with prior studies that have highlighted the differential expression of GAG sulfation enzymes and core protein expression across diverse cell types, with conserved expression of the initiation and elongation enzymatic machinery [54, 55].

Importantly, transcriptomic profiling and qPCR analysis of HEK293T cells confirmed that *HS3ST1* and *HS3ST5* are expressed at low but detectable basal levels (Fig. 1E, Suppl. Fig. 1A). Consistent with this expression profile, AT is known to interact with HS through both high-affinity pentasaccharide-dependent binding, in which 3-*O* sulfation is an essential structural determinant, and lower-affinity charge-dependent interactions driven by overall sulfation density [56, 57], and both modes are likely operative in our AT binding experiments. The selective enhancement of AT binding upon *SDC1* activation therefore likely reflects both the elevated 6-*O*-sulfation density on SDC1-associated HS chains and their intrinsically superior capacity to serve as 3-*O*-sulfotransferase substrates [58]. This is also directly supported by cell-free HS3ST1/5 sulfation assays and by the amplified AT binding response in vascular endothelial cells (HUVECs), where *HS3ST1* is endogenously expressed [59, 60]. Interestingly, our CRISPRa screens also uncovered shared enrichment of *GPC1* and *HS3ST4* across both screens (Fig. 2), suggesting that additional proteoglycans or biosynthetic enzymes may also influence AT binding and/or total HS abundance. In fact, a recent study showed that overexpression of *HS3ST4* in CHO cells led to enhanced HS 3-*O* sulfation and anticoagulant activity [61]. In addition, *PTGER2* emerged as a shared hit in both screens despite not previously being implicated in HSPG assembly. *PTGER2* encodes the prostaglandin E2 receptor EP2, a Gs-coupled GPCR that activates cAMP/PKA-dependent signaling and downstream transcriptional programs through CREB [62]. Although the mechanism remains to be determined, PTGER2-mediated signaling could influence HS assembly through downstream effects on gene expression, enzyme activity, and/or secretory pathway dynamics. Future work will investigate the role of these factors in the regulation and production of anticoagulant HS.

The dichotomy we observed between SDC1 and SDC2 within the syndecan family raises important questions about proteoglycan assembly and structure/function relationships. Both syndecans are type I transmembrane proteoglycans that share conserved transmembrane and cytoplasmic domains, yet their ectodomains differ substantially in length and amino acid sequence and they are differentially expressed across tissues [37]. In adult tissues, SDC1 is predominantly expressed by epithelial cells and plasma cells, with particularly high levels in squamous epithelia, gastrointestinal mucosa, and hepatocytes [63, 64]. Conversely, SDC2 is the predominant syndecan of mesenchymal and stromal compartments, with high expression in fibroblasts and endothelial cells [65]. This reciprocal distribution raises the possibility that the cell type-specific expression of proteoglycans may be one mechanism by which tissues regulate the sulfation complexity of their surface HS. Consistent with their distinct expression patterns, studies of systemic *Sdc1/Sdc2* knockout mouse models show non-overlapping phenotypes, with *Sdc1* deficiency driving metabolic dysregulation and altered lipid clearance [66, 67] while *Sdc2* deletion selectively disrupts developmental angiogenesis and hematopoietic stem cell quiescence [68, 69]. These divergent phenotypes provide evidence that syndecans serve functionally distinct biological roles that cannot be attributed solely to differences in expression level or tissue distribution but likely also reflect differences in their GAG composition and binding partners. Our findings support this interpretation and are consistent with prior work demonstrating that the N-terminal ectodomain sequence of SDC2 selectively promotes 6-*O*-sulfation of its HS chains relative to SDC4, driving VEGFA_165_-dependent angiogenesis [68].

Structural analyses of GAGs isolated from recombinant ectodomains of SDC1 and SDC2 provided direct biochemical evidence for core protein-dependent differences in GAG sulfation patterning. eSDC1-derived HS contained higher levels of 6-*O* sulfated disaccharides across multiple species as well as elevated levels of critical 3-*O*-sulfated tetrasaccharide sequences consistent with enrichment of AT-binding sites (Fig. 5D). Moreover, CS chains were more highly sulfated in SDC1-derived material (Fig. 5E). Altogether, these data indicate that the ectodomain of the core protein, which is responsible for most of the sequence differences within the syndecan family, influences the sulfation patterning of its attached GAG chains. Using ectodomain-derived HS in 3-*O* sulfation assays, we demonstrated that eSDC1-HS served as a better substrate for HS3ST1 and HS3ST5, which we propose is likely driven by the elevated 6-*O* sulfated sites that are a prerequisite for AT-binding site formation [58, 70]. These results connect the capacity for 3-*O* sulfation to not only the expression level of 3-*O* sulfotransferases, but also to the upstream sulfation context of the HS chain itself, which we propose is influenced by core protein identity.

The molecular basis for the observed differential cell surface trafficking kinetics between SDC1 and SDC2 remains an open and important question. The difference in cell surface recovery could arise from varied biosynthetic rates and/or Golgi residence times, with SDC1 transiting the Golgi longer, allowing for more time for HS chains to be modified. This Golgi residence time model is mechanistically attractive because it connects our kinetic observation directly to the HS sulfation patterning data without invoking direct communication between sulfotransferase enzymes and the core protein sequence. The observation that cycloheximide treatment fully blocked SDC1/SDC2 recovery (Fig. 6B) points to a lack of difference in recycling rates and/or utilization of pre-formed intracellular pools. In addition, ectodomain length and topology may also influence the efficiency of vesicle packaging and/or interactions with cargo receptors during anterograde transport. SDC1 possesses a substantially longer ectodomain than SDC2, and there is precedent for ectodomain size and other physicochemical properties, including glycosylation, influencing Golgi transit kinetics for other transmembrane proteins [71, 72]. Although the syndecan cytoplasmic tails are highly conserved at the sequence level, SDC1 and SDC2 have been shown to interact differentially with PDZ domain-containing scaffolds, which could influence proteoglycan localization and recycling at the plasma membrane [73].

Finally, the number and occupancy of GAG attachment sites, which are known to vary among syndecan family members [74], may itself function to regulate trafficking and export competence. Distinguishing among these possibilities will require direct measurement of Golgi transit times using pulse-chase or organelle-specific proximity labeling approaches in future work. Regardless of the precise step that differs, the slower overall biosynthetic flux of SDC1 is consistent with a model in which the core protein indirectly shapes HS modification outcomes through differences in the kinetics of its transit through the biosynthetic pathway.

More broadly, this work highlights the importance of considering proteoglycan core protein identity when interpreting GAG function in any biological context. HSPGs are differentially expressed across tissues and cell types, and this has generally been interpreted through the lens of extracellular domain interactions and co-receptor function. Our findings suggest that the choice of core protein expressed by a given cell type may systematically regulate the sulfation patterning of its cell surface HS and subsequent HS-ligand interactions at the cell surface and ECM interface. This principle may be particularly relevant in disease contexts where proteoglycan expression profiles are altered. SDC2 upregulation and its resultant HS sulfation patterning impacts hematopoietic stem cell self-renewal [75] and can regulate the immune response during sepsis [76]. In certain cancers, proteoglycan expression profiles are distinctly altered and are implicated in tumorigenesis. For example, the marked upregulation of SDC1 that occurs in multiple myeloma and certain epithelial cancers may influence HS sulfation in ways that affect growth factor signaling and tumor microenvironment organization beyond simply increasing HSPG abundance [77–79].

Ultimately, the screening framework established here, pairing genome-wide CRISPRa perturbation with functional HS-ligand binding readouts, provides a scalable platform for dissecting the determinants of HS functional specificity across diverse biological contexts, and motivates systematic investigation of how proteoglycan core protein selection shapes the HS code in development, homeostasis, and disease.

## Materials and Methods

### Cell Culture

HEK293T cells (ATCC, CRL-3216) were grown in DMEM (Gibco) supplemented with 10% (v/v) FBS (Cytiva) and 1% (v/v) penicillin/streptomycin (Gibco) at 37°C under an atmosphere of 5% CO_2_/95% air. HUVECs (Lonza) were grown in EGM-2 media supplemented with the EGM-2 BulletKit (Lonza) on tissue culture plates coated with 0.2% gelatin. All cell lines were passaged every 2-4 days and revived from liquid nitrogen after ≤10 passages. HUVECs were used between 2-6 passages.

### Virus Production and Cell Line Generation

HEK293T cells were co-transfected with FuGENE 6 (Promega) and each of the following plasmids at 4 µg each: viral envelope plasmid (pMD2.g, Addgene #12259, gift from Didier Trono), packaging plasmid (psPAX2, Addgene, #12260, gift from Didier Trono), and dCas9-VP64 expression plasmid (lenti dCas9-VP64_Blast, Addgene #61425, gift from Feng Zhang). Lentiviral particles were collected and added to HEK293T cells followed by selection with 10 µg/ml blasticidin for 4 days to generate stable dCas9-expressing cells. For the SAM CRISPRa system [39], HEK293T-dCas9-MPHv2 cells were generated in a similar fashion by producing lentiviral particles containing the lentiMPHv2 plasmid (Addgene #89308, gift from Feng Zhang) in HEK293T cells and subsequent transduction of dCas9-expressing HEK293T cells and selection with 400 µg/ml hygromycin for 4 days. Individual sgRNA CRISPRa cell lines were generated by ligating sgRNA targeting sequences into either pXPR_502 (Addgene #96923, gift from John Doench and David Root) or lenti sgRNA(MS2)_puro optimized (Addgene #73797, gift from Feng Zhang). Lentiviral particles were produced by co-transfection of the sgRNA plasmid with pMD2.G and psPAX2 into HEK293T cells using FuGENE 6, collected, and used to transduce either HEK293T-dCas9 or HEK293T-dCas9-MPHv2 cells. After selection of the transduced cells with puromycin (0.5 µg/ml) for 3 days, the surviving cells were used in experiments as stable polyclonal populations. A list of sgRNA sequences are provided in Supplementary Table 5.

### Western blotting

Total protein was extracted from cells using RIPA Lysis and Extraction Buffer (EMD Millipore) supplemented with protease inhibitors (Roche). Protein concentration was determined by BCA assay (Thermo Scientific-Pierce). Protein samples were subjected to SDS-polyacrylamide gel electrophoresis (4-12% Bis-Tris, Invitrogen), blotted on nitrocellulose membranes (Invitrogen), and probed for Cas9 (mouse IgG anti-Cas9, Thermo Fisher #MA1-201, 1:500) and GAPDH (rabbit IgG anti-GAPDH, Cell Signaling Technology #5174, 1:2000). Membranes were blocked with 5% milk in Tris-buffered saline with 0.1% Tween (TBST) for 1 hour at room temperature then incubated with the respective primary antibodies at 4°C overnight. Membranes were then incubated with IRDye secondary antibodies (1:14,000; LI-COR Biosciences) matched to the primary antibody host species and visualized using an Odyssey imaging system (LI-COR Biosciences). Protein sizes were estimated using a protein ladder (Thermo Fisher, #26616). Uncropped western blot source data is included in Suppl. Fig. 4.

### Transfection Experiments

HUVEC cells were seeded at a density of 1.5 x 10^5^ cells per well in 6-well tissue culture plates pre-coated with 0.2% gelatin. 24 hours after seeding, cells were washed with PBS and transfected with Lipofectamine LTX with Plus reagent (Invitrogen) and 100 ng of human N-FLAG-SDC1 expression plasmid (Sino Biological, #HG11429-NF) or human N-FLAG-SDC2 expression plasmid (Sino Biological, #HG10355-NF) in 1 ml of serum-free EGM-2 basal medium according to the manufacturer’s instructions. Cells were incubated with this mixture for 4 hours, after which the medium was replaced with complete EGM-2 media. Flow cytometry experiments were performed 48 hours post-transfection.

### Lentiviral Library Production and Titer Determination

The pooled human Calabrese P65-HSF library (set A) was obtained from Addgene (Addgene #92379, gift from David Root and John Doench), transformed and amplified in ElectroMAX Stbl4 Competent cells (Invitrogen, #11-635-018) via electroporation (Gene Pulser XCell, Bio-Rad), and purified using the ZymoPURE II Plasmid Maxiprep Kit (Zymo Research). Next-generation sequencing (MiSeq) of the amplified plasmid library confirmed coverage of >98% of sgRNAs and 100% of genes represented in the Calabrese library. 24 hours prior to transfection, 1.8 x 10^7^ HEK293T wild-type cells were seeded into four T175 flasks (175 cm^3^). The next day, the cells were washed twice with PBS, and 25 ml of DMEM + 10% FBS (without antibiotics) was added to each flask. In 15 ml tubes, a mixture of 40 µg of the Calabrese plasmid library, 5 µg of lentiviral envelope plasmid (pMD2.g; Addgene, #12259) and 50 µg of lentiviral packaging plasmid (psPAX2; Addgene, #12260) were diluted in 6 ml of OptiMEM and supplemented with 305 µL of TransIT-LT1 transfection reagent (Mirus Bio, #MIR2300). The mixture was incubated at room temperature for 30 minutes and then added dropwise to each flask, respectively, with gentle shaking. After an 8-hour incubation at 37°C, the transfection mix was replaced with 35 ml of DMEM +10% FBS + 1% BSA and the cells were incubated for an additional 36-hour incubation. Subsequently, the virus-containing medium was collected, centrifuged for 10 min at 3,000 x *g* (4°C), and filtered through a 0.45-μm PES syringe filter (Thermo Scientific, #09-740-114). The viral stock was aliquoted, snap frozen, and stored at -80°C.

To determine lentiviral titer, HEK293T-dCas9 cells were transduced in 12-well plates via spinfection with a range of virus volumes with 3.0 x 10^6^ cells per well in the presence of 8 µg/ml polybrene. The plates were centrifuged at 2,000 rpm for 2 hours at 37°C and were then transferred to a 37°C incubator for 4 hours. Each well was then trypsinized, and an equal number of cells seeded into each of two wells of a 96-well plate. Two days post-transduction, puromycin was added to one well out of the pair. After 3 days, both wells were assessed for viability using the Cell Titer Blue assay (Promega). A viral dose corresponding to an MOI of ∼0.3 was used for subsequent library screening. MOI was calculated using the formula: µ = -log (1 - q), where q is the fraction of cells surviving puromycin selection.

### CRISPRa Library Transduction and Screening Assays

HEK293T-dCas9 cells were transduced with the Calabrese library at a low MOI (∼0.3). Transductions were performed with enough cells to achieve a representation of at least 500 cells per sgRNA, taking into account a ∼30% transduction efficiency. A total of 10.8 x 10^7^ HEK293T-dCas9 cells were distributed across three 12-well plates in complete DMEM medium containing 8 µg/ml polybrene (Sigma) and the virus suspension. Plates were centrifuged at 2,000 rpm for 2 hours (37°C) with the viral containing media, after which the media was swapped with fresh complete DMEM media, and cells were incubated overnight. The next day, cells were lifted with trypsin, combined, then split evenly into thirty 15 cm plates and incubated with medium containing puromycin (0.5 µg/ml) for 6 days. Aliquots of the selected Calabrese cell library were frozen and stored in liquid nitrogen.

For the FACS-based screening assays, Calabrese library-transduced cells (4 x 10^7^ cells per replicate) were thawed and cultured across multiple 15 cm plates for 2 days. On the day of sorting, cells were washed with PBS, gently lifted with PBS (+ 10 mM EDTA), and transferred to 50 ml conical tubes. For binding assays, cells were incubated in a suspension of PBS + 0.1% BSA with either 0.5 µg/ml mAb 10E4 (AMSBio, #370255-1, Clone F58-10E4, 1:1,000) for 30 min at 4°C or with 500 nM human antithrombin (Aniara Diagnostica, #APP004B) for 1 hour at 4°C. After primary incubation, cells were washed twice with PBS + 0.1% BSA prior to incubation with appropriate secondary antibodies. Bound 10E4 was detected with 2 µg/ml anti-mouse IgM conjugated to AlexaFluor647 (Invitrogen, # A-21238, 1:1,000), incubated for 15 min at 4°C. Bound antithrombin was detected with 2 µg/ml anti-AT pAb (R&D Systems, #AF1267, 1:100), incubated for 30 min at 4°C, followed by 2.5 µg/ml donkey anti-goat IgG conjugated to AlexaFluor488 (Invitrogen, #A-11055, 1:1,000), incubated for 15 min at 4°C. Each cell population was collected, resuspended in PBS + 0.1% BSA (+ 10 mM EDTA), and FACS sorted for the top 1% 10E4+ or AT+ cells on a MoFlo Astrios cell sorter (Beckman Coulter) at the UGA CTEGD Cytometry Shared Resource Laboratory. Each sorted cell population was pelleted by centrifugation (4°C), the supernatant was removed, and cell pellets were snap frozen and stored at -80°C for genomic DNA isolation.

### Genomic DNA Preparation and Sequencing

Genomic DNA of sorted cell populations was extracted using the DNeasy Blood & Tissue Kit (Qiagen, #69506), and the stably integrated sgRNAs were amplified through PCR. Genomic DNA extracts were evenly divided into multiple PCR reactions using ∼8-10 µg of DNA per reaction. Per reaction tube, genomic DNA was combined with 1.5 µL of ExTaq polymerase (TaKaRa), 8 µL of 10 mM deoxynucleotide triphosphate (dNTP; TaKaRa), 0.5 µL of 100 µM of P5 primer mix, 10 µL of 5 µM P7 primer (with barcode), 10 µL of 10X reaction buffer (TaKaRa), and UltraPure water (Invitrogen) for a final reaction volume of 100 µL. PCR products were generated with the following reaction conditions: 95°C preheat, 1 min at 95°C, 30 sec at 95°C for denaturing, 30 sec 53°C for annealing, and 30 sec at 72°C for extension, for 26-28 cycles before a 10 min hold at 72°C. Upon completion, 20 µL from each PCR reaction was pooled per sample and purified using AMPure XP beads (Beckman Coulter, #A63880) according to the manufacturer’s instructions. Purified PCR products were combined per sample and the concentration was determined using Qubit with the dsDNA high-sensitivity (HS) assay (Thermo Fisher). All samples were pooled and amplicons were sequenced on a NovaSeq6000 (Illumina).

### Screening Sequencing Analysis

Analysis of the raw amplicon sequencing data, including quality control, sequence alignment, read counting, sgRNA enrichment analysis, and gene ranking, was carried out using PinAPL-Py v2.9.2 [35]. Sequencing reads were trimmed using CutAdapt to remove the 5’ constant region (5’-TCTTGTGGAAAGGACGAAACACCN-3’), retaining reads with a minimum length of 20 nucleotides. Trimmed reads were aligned to the Calabrese sgRNA library using Bowtie2 with the following parameters: seed length of 11, one seed mismatch permitted, score function S,1,0.75, and a minimum alignment score of 40. A cut error tolerance of 0.1 was applied, and alignment files were removed after read counting. Per-sgRNA read counts were normalized by counts per million (CPM), and replicate samples were averaged using the median. sgRNA-level statistical analysis was performed using a two-sided test with Benjamini-Hochberg false discovery rate correction (α = 0.01), with sgRNAs clustered by counts. Gene-level ranking was performed using the SigmaLFC metric with 10,000 permutations (α = 0.05). Gene ranking scores, p-values, and number of significant sgRNAs per gene (minimum 2) were used to rank and prioritize gene hits for downstream validation.

### Flow Cytometry Experiments

HEK293T cells grown in monolayer culture were washed with PBS (Gibco) and lifted using PBS with 10 mM EDTA. HUVEC cells were lifted with Accutase (Sigma-Aldrich). Cells were counted and 100,000 cells were seeded into each well of a 96-well round bottom plate (Corning). For binding assays, cells were incubated in a suspension of PBS + 0.1% BSA for 30 min at 4°C with 0.5 µg/ml mAb 10E4 (AMSBio, #370255-1, Clone F58-10E4, 1:1,000), 1 µg/ml mAb 3G10 (AMSBio, #370260, clone F69-3G10, 1:1000), 1.4 µg/ml mAb CS56 (Sigma-Aldrich, #C8035, clone CS56, 1:1000), mAb anti-FLAG (Sigma-Aldrich, #F3165, 1:1000), 80 nM biotin-FGF1 (Peprotech, #100-17A), and 2.5 nM biotin-FGF2 (Peprotech, #100-18B). To enable detection of cell surface syndecans, cells were pre-treated with a cocktail of heparin lyases I, II, and III (5 mU/mL each; IBEX) and chondroitinase ABC (20 mU, Sigma-Aldrich, #C3667) in PBS + 0.1% BSA for 30 minutes at 37°C prior to incubation with anti-syndecan primary antibodies.

Recombinant proteins were biotinylated in-house, as previously described [80]. Alternatively, cells were incubated for 1 hour at 4°C with 500 nM human antithrombin (Aniara), 2 µg/ml mAb Anti-SDC1 (Novus, #NB100-64980, clone B-A38), 2 µg/ml pAb Anti-SDC2 (R&D Systems, #AF2965), VEGF_165_ (Peprotech, #100-20, 1:2000). After primary incubation, cells were washed twice with PBS + 0.1% BSA followed by incubation with appropriate secondary antibodies.

Bound antithrombin was detected with 2 µg/ml anti-AT pAb (R&D Systems, #AF1267, 1:100), incubated for 30 min at 4°C, followed by 2.5 µg/ml donkey anti-goat IgG conjugated to AlexaFluor488 (Invitrogen, #A-11055, 1:1000), incubated for 15 min at 4°C. Bound VEGF_165_ was detected with pAb biotin-VEGF_165_ (PeproTech, #500-P10BT-25UG), incubated for 30 min at 4°C. Binding of biotinylated proteins was detected by streptavidin-Cy5 (Molecular Probes, #434316, 1:1000), incubated for 30 min at 4°C. All other fluorescence anti-bodies used were diluted 1:1000 and incubated for 15 min on at 4°C: goat anti-mouse IgM conjugated to AlexaFluor647 (Thermo, #A21238) and goat anti-mouse IgG conjugated to AlexaFluor647 (Fisher, #A21235). For heparin lyase pretreatment experiments, adherent cells were incubated with 5 mU/ml each of heparin lyases I, II and III (IBEX) for 30 min at 37°C in PBS + 0.1% BSA. Flow cytometry analysis was performed on a CytoFlex S (Beckman Coulter) instrument (≥ 10,000 events per sample) and raw data were analyzed using FlowJo Analytical Software (Tree Star, Inc.). Cells were gated according to forward and side scattering (Suppl. Fig. 3). The extent of protein binding was quantified using the geometric mean of the fluorescence intensity. These values were plotted and further analyzed using GraphPad Prism v.8.0.

### RNA Isolation, cDNA Synthesis, and Quantitative PCR

RNA was isolated from cell lines and transfected cells using TRIzol (Invitrogen) and the Direct-zol RNA Miniprep kit (Zymo) following the manufacturer’s instructions. cDNA was prepared from total RNA using the SuperScript IV First Strand Synthesis kit (Invitrogen) using random hexamers following the manufacturer’s instructions. Quantitative PCR (qPCR) was performed using cDNA and SYBR Green Master Mix (Applied Biosystems) following the manufacturer’s instructions. The expression of *YWHAZ* (housekeeping gene) was used to normalize the expression of target genes between samples. The primers used for quantitative PCR are provided in Supplementary Table 6.

### Recombinant Expression of Syndecan-1 and Syndecan-2 Ectodomains

The expression constructs were generated encoding the truncated ectodomain of human Syndecan-1 (UniProt P18827, residues 23-251) and human Syndecan-2 (UniProt P34741, residues 19-144) as an NH_2_-terminal fusion protein in the pGEn2 expression vector essentially as described in prior studies [43]. Briefly, the fusion protein coding region was comprised of a 25-amino acid signal sequence, an His_8_ tag, AviTag, the “superfolder” GFP coding region, the 7-amino acid recognition sequence of the tobacco etch virus (TEV) protease followed by the truncated Syndecan-1 or Syndecan-2 coding regions. The Syndecan-1 and Syndecan-2 expression constructs were used for transient transfection of suspension culture HEK293-F cells (FreeStyle™ 293-F cells, Thermo Scientific) maintained at 0.5-3.0x10^6^ cells/ml in a humidified CO_2_ platform shaker incubator at 37°C with 50% relative humidity and 125 rpm. Transient transfection was performed using HEK293-F cells in expression medium comprised of a 9:1 ratio of Freestyle™ 293 expression medium (Thermo Scientific) and EX-Cell expression medium including Glutamax (Sigma-Aldrich). Transfection was initiated by the addition of plasmid DNA and polyethyleneimine as transfection reagent (linear 25-kDa polyethyleneimine, Polysciences, Inc.). 24 hours post-transfection, the cell cultures were diluted with an equal volume of fresh media supplemented with valproic acid (2.2 mM final concentration) and protein production was continued for an additional 5 days at 37°C and 125 rpm. The cell cultures were harvested, clarified by sequential centrifugation at 1200 rpm for 10 min and 3500 rpm for 15 min at 4°C, and passed through a 0.8 µM filter (Millipore). The protein preparations were adjusted to contain 20 mM HEPES, 20 mM imidazole, 300 mM NaCl, pH 7.5, and subjected to Ni-NTA Superflow (Qiagen) chromatography using a column preequilibrated with 20 mM HEPES, 300 mM NaCl, 20 mM imidazole, pH 7.5 (Buffer I). Following loading of the sample, the column was washed with 3 column volumes of Buffer I followed by 3 column volumes of Buffer I containing 50 mM imidazole and eluted with Buffer I containing 300 mM imidazole at pH 7.0. The protein was concentrated to 1 mg/ml using an ultrafiltration pressure cell (Millipore) with a 10-kDa molecular mass cutoff membrane and buffer exchanged with 20 mM HEPES, 150 mM NaCl, pH 7.0, 10% glycerol. The final protein preparations were aliquoted and stored at -80°C until use.

### Glycosaminoglycan Isolation and Purification

GAG chains were released from recombinant syndecan ectodomains through β-elimination by adding NaOH to protein samples to a final concentration of 0.4 M, as previously described [81]. Subsequently, samples were incubated overnight at 4°C then neutralized with an equal volume of 0.4 M glacial acetic acid. Neutralized samples were diluted 1:20 in wash buffer (200 mM NaCl, 50 mM sodium acetate, pH 6.0) and then purified as previously described [81]. Briefly, a 0.5 ml bed of DEAE-Sephacel (Cytiva) was prepared using equilibration buffer (200 mM NaCl, 50 mM sodium acetate, 0.1% Triton-X-100, pH 6.0) in Poly-Prep Columns (0.8 x 4 cm, 10 mL volume, Bio-Rad). Samples were loaded onto the column, washed twice with wash buffer, and eluted with 1.5x bed volume of elution buffer (2 M NaCl, 50 mM sodium acetate, 6.0 pH). Eluates were then desalted using PD-10 desalting columns with Sephadex G-25 resin (Cytiva). Desalted samples were lyophilized overnight and stored at -80°C.

### LC-MS Analysis of HS Disaccharides

For HS disaccharide analysis, GAGs isolated from eSDC1 and eSDC2 were digested with 2 mU each of heparin lyases I, II and III (IBEX) in 40OmM ammonium acetate, 3.3 mM calcium acetate (pH 7.0), at 37°C for 16 hours. Disaccharides were aniline-tagged and analyzed by HILIC-Q-TOF-MS on the Waters SYNAPT XS mass spectrometer equipped with an ACQUITY UHPLC H-class system (BEH glycan column, 2.1Omm x 100 mm), as previously described [45]. Mobile phase A was 50OmM ammonium formate (pH 4.4); B was 100% acetonitrile. The elution was as follows: 0-5.0 min with isocratic 10% A, linear gradient 10-33% A from 5.0 to 49.0Omin, linear gradient 33-45% A from 49.0 to 51.5 min, isocratic 45% A from 52.0 to 60 min, then the column was washed with 90% B for 10 min. Column temperature was at room temperature; the flow rate was 0.5 mL min^−1^, and the injection volume was 2 µL. The total HS content was normalized to input protein amounts.

### Sample Preparation of HS Tetrasaccharides and CS/DS Disaccharides

For HS tetrasaccharide and CS/DS disaccharide analysis, intact eSDC1 and eSDC2 protein were used for GAG isolation and were analyzed, as previously described [44]. Proteins were subjected to a 3kDa MWCO spin column (MilliporeSigma, #UFC500324) and centrifuged at 15,000 rpm for 10 minutes at room temperature. The retentate was kept in the 3kDa MWCO column and washed three times with 200 µL of deionized water at 15,000 rpm for 10 minutes at room temperature to get rid of salt in the sample solution. After the desalting procedure, the retentate on the membrane was recovered, 500 ng each of ^13^C-labeled 3-*O*-sulfated oligosaccharide calibrants were added to the solution, and 100 µL of heparin lyase and chondroitinase ABC digestion solution were added to the retentate-calibrant mixture. The digestion solution contained 7.5 µL enzymatic buffer (100 mM sodium acetate, 2 mM calcium acetate buffer (pH 7.0) containing 0.1 g/L BSA), 1.25 µL heparin lyase I (2.49 mg/mL), 2.5 µL heparin lyase II (13.6 mg/mL), 3 µL chondroitinase ABC (6 mg/mL), and 85.75 µL H_2_O. The reaction solution was incubated overnight at 37°C. Before recovering the digests from the digest solution, a known amount ^13^C-labeled CS disaccharide calibrants were added to the digestion solution. The HS tetrasaccharides and CS/DS disaccharides were recovered by 30kDa MWCO column (15,000 rpm for 10 minutes at room temperature), and the filter unit was washed twice with 200 µL of deionized water (15,000 rpm for 10 minutes at room temperature). The collected filtrates were freeze-dried before the 2-aminoacridone (AMAC) derivatization.

A total of five ^13^C-labeled 3-*O*-sulfated oligosaccharide calibrants were used during the analysis of 3-*O*-sulfated tetrasaccharides. These structures are oligo 1: GlcNAc-GlcA-GlcNAc6S-GlcA*-GlcNS3S6S-IdoA2S*-GlcNS6S-GlcA-pNP; oligo 2: GlcNAc-GlcA-GlcNS6S-GlcA*-GlcNS3S6S-IdoA2S-GlcNS6S-GlcA-pNP; oligo 3: GlcNS6S-GlcA*-GlcNS6S-IdoA2S*-GlcNS3S6S-IdoA2S*-GlcNS6S-GlcA-pNP; oligo 4: GlcNS-GlcA-GlcNS-IdoA2S*-GlcNS3S-IdoA2S-GlcNS-GlcA-pNP; and oligo 5: GlcNAc-GlcA-GlcNS-IdoA2S*-GlcNS-IdoA2S-GlcNS3S-IdoA2S-GlcNS-GlcA-pNP. Four ^13^C-labeled CS disaccharide calibrants were used for the analysis of CS portion: ΔUA*-GalNAc, ΔUA*-GalNAc4S, ΔUA*-GalNAc6S, ΔUA*-GalNAc4S6S.

### Chemical Derivatization and LC-MS/MS Analysis of Disaccharides and Tetrasaccharides

The AMAC (#06627, Sigma-Aldrich) derivatization of lyophilized samples was performed by adding 5 μL of 0.1 M AMAC solution in DMSO/glacial acetic acid (17:3, v/v) and incubating at room temperature for 10 minutes. Then 5 μL of freshly prepared 1 M aqueous sodium cyanoborohydride (Thermo Fisher, #087839-06) was added to this solution. The reaction mixture was incubated at 45°C for 2 hours. After incubation, the reaction solution was centrifuged (15,000 rpm for 10 minutes at room temperature) to obtain the supernatant that was subjected to the LC-MS/MS analysis.

The analysis of AMAC-labeled HS and CS was performed on a Vanquish Flex UHPLC System (Thermo Fisher) coupled with TSQ Fortis triple-quadrupole mass spectrometry as the detector. The ACQUITY Glycan BEH Amide column (1.7 µm, 2.1 x 150 mm; Waters, Ireland, UK) was used to separate tetrasaccharides at 60°C. Mobile phase A was 50 mM ammonium formate in water, pH 4.4. Mobile phase B was acetonitrile. The elution gradient is as follows: 0-15 minutes 83-70% B, 15-30 minutes 70-50% B, 30-35 minutes 50% B, 35-45 minutes 83% B. The flow rate was 0.3 mL/minute. On-line triple-quadrupole mass spectrometry operating in the multiple reaction monitoring mode was used as the detector. The electrospray ionization-mass spectrometry analysis was operated in the negative-ion mode using the following parameters:

Neg ion spray voltage at 3.0 kV, sheath gas at 55 Arb, aux gas 25 arb, ion transfer tube temp at 250°C and vaporizer temp at 400°C. TraceFinder software was applied for data processing. The amount of HS and CS was determined by comparing the peak area of native di/tetra-saccharides to each di-/tetrasaccharide calibrant.

### Expression and Purification of Recombinant Enzymes

Transmembrane region-truncated cDNAs encoding *Homo sapiens* HS3ST1 (UniProt O14792) and HS3ST5 (Q8IZT8), *Pasteurella multocida* inorganic pyrophosphatase (PmPpA, P57918), *Saccharomyces cerevisiae* sulfate adenylyltransferase (ScATPS, P08536), and *Escherichia coli* adenylyl-sulfate kinase (EcAPSK, P0A6J1) were obtained from the Mammalian Gene Collection or synthesized de novo (Twist Bioscience). Human enzyme genes were subcloned into the pGEn2 vector whereas bacterial enzyme genes were subcloned into pET21a or pET15b expression vectors containing N-terminal green fluorescence protein fusions.

Bacterial expression plasmids were transformed into *E. coli* BL21 (NEB) or ArticExpress (Agilent) competent cells by heat shock. Single colonies were inoculated into Luria-Bertani broth and cultured at 37°C until OD_600_ reached 0.6. Cultures were then cooled to 16°C and induced with 0.5 mM isopropyl-β-D-1-thiogalactopyranoside (GoldBio) for 16 hours. Mammalian expression plasmids were transfected into FreeStyle293F (Thermo Fisher), as previously described [43]. Bacterial cell pellets were lysed by sonication, and mammalian cell supernatants were collected by centrifugation. Recombinant proteins were purified using Ni-NTA Superflow affinity chromatography, as previously described[43]. Imidazole was removed by repeated ultrafiltration. Protein purity was assessed by gel electrophoresis.

### [^35^S]PAPS Generation

Radiolabeled 3’-phosphoadenosine-5’-phosphosulfate (PAPS) was produced as described previously [82]. Briefly, [^35^S]PAPS was generated in reaction quantities of 100 µL reaction volumes under the following buffered conditions: 10 mM MgCl_2_, 10 mM LiCl, 50 mM Tris, 8.0 pH. For the reaction,130 µg of adenosine 5’-triphosphate sulfurylase (ATPS) (from *K. lactis*), 1 mg of adenosine 5’-phosphosulfate kinase (APSK) (from *Penicillium chrysogenum*), and 600 µg of pyrophosphatase (PpA) (from *E. coli*) were combined with 10 mM ATP, 10 million cpm of sodium [^35^S]sulfate (#NEX041001MC, Revvity Health Sciences Inc.), 10X reaction buffer, and pure water to a total volume of 100 µL. The reaction was then incubated at 37°C for 6 hours. Crude [^35^S]PAPS was stored at -80°C until use.

### In Vitro 3-O Sulfation Reactions

All sulfation reactions were performed under the following buffer conditions: 5 mM MnCl_2_, 5 mM MgCl_2_, 2.5 mM CaCl_2_, 50 mM MES, 120 µg/ml BSA, 20 µg/ml protamine chloride, 0.5% Triton-X, pH 7.0. Importantly, MnCl_2_ was added after pH adjustment to prevent precipitation at alkaline pH. For *in vitro* sulfation reactions, varying amounts of HS3ST1 or HS3ST5 enzyme were added to a fixed quantity of HS isolated from eSDC1 or eSDC2. Equal mass amounts of eSDC1 and eSDC2 HS were loaded into each respective reaction based on total HS abundance quantified by disaccharide analysis, as outlined above. Sulfation reactions were carried out in reaction volumes of 50 µL with 500,000 cpm of [^35^S]PAPS. Reactions were incubated for 18 hours at 37°C. For downstream analyses of 3-*O*-sulfated HS chains, reaction mixtures were diluted 1:20 in wash buffer (250 mM NaCl, 50 mM sodium acetate, pH 6.0) and purified over DEAE-Sephacel (250 µL bed volume, Cytiva), as described above. DEAE columns were washed twice with wash buffer to remove unreacted [^35^S]PAPS and free [^35^S]sulfate, and [^35^S]-labeled chains were released with elution buffer (2 M NaCl, 50 mM sodium acetate, pH 6.0) and a portion was used to quantify [^35^S] incorporation with a liquid scintillation counter (Beckman Coulter, LS6500).

### Size Exclusion Chromatography of [^35^S]-labeled Heparan Sulfate

[^35^S]-labeled HS chains isolated from eSDC1 or eSDC2 (300 ng) were 3-*O*-sulfated using 70 ng HS3ST1 enzyme and purified, as described above. Subsequently, 3-*O*-sulfated [^35^S]-labeled HS chains were analyzed by size exclusion chromatography (Sepharose CL-6B column 1.7 cm x 80 cm; 50mM sodium acetate, 0.2M NaCl, pH 6.0). The average molecular weight of HS chains was determined based on previous size determinations using a Sepharose 6B column [83].

### Cell Surface Recovery Assays

To assess cell surface recovery kinetics of SDC1 and SDC2, HEK293T wild-type cells were seeded at a density of 1.5 x 10^5^ cells per well in 6-well tissue culture plates 48 hours prior to the experiment. On the day of the assay, baseline cell surface levels of SDC1 and SDC2 were established by gently detaching cells with PBS supplemented with 10 mM EDTA, followed by immediate collection and flow cytometry analysis, as described above. To remove surface-exposed proteins, parallel cultures were incubated with 0.25% trypsin-EDTA (Gibco) for 5 minutes at 37°C. Trypsin was immediately neutralized by addition of an equal volume of complete DMEM containing 10% FBS, and cells were spun down and washed twice with PBS. Trypsin-released cells were either collected on ice to establish a trypsin baseline or replated at equal density into 6-well tissue culture plates in complete DMEM and incubated at 37°C to allow surface protein recovery. At 2, 4, 6, or 8 hours post-replating, cells were lifted with PBS supplemented with 10 mM EDTA, collected, and analyzed by flow cytometry for SDC1 and SDC2 surface levels using anti-SDC1 and anti-SDC2 antibodies, as described above. Cell surface recovery at each time point was expressed as a percentage of the EDTA-established baseline signal for each respective proteoglycan. Initial rates of surface recovery were calculated from the slope of the linear fit of the recovery curve over the 8-hour time course.

For cycloheximide experiments, cells were pre-treated with cycloheximide (10 µg/ml, Thermo Fisher, #J6690103) for 1 hour prior to trypsinization. Cells were either collected on ice or replated with media containing cycloheximide throughout the entire 0-8 hour recovery period. Cell surface recovery of SDC1 and SDC2 was then assessed at the same time points described above. All flow cytometry data were collected on a CytoFlex S instrument (Beckman Coulter) and analyzed using FlowJo software (Tree Star, Inc.).

### Statistics and reproducibility

Statistical tests, sample sizes, and numbers of biological replicates are indicated in the figure legends. Statistical significance was indicated by the following: *p<0.05, **p<0.01, ***p<0.001, ****p<0.0001. All tests were two-tailed and performed using GraphPad Prism v.8.0 or v10.6 software. Measurements were taken from distinct biological samples, and the number of biological replicates is indicated in figure legends. All error bars represent mean ± standard deviation. Western blots were performed three times independently and representative images are shown in the figures.

## Data availability

Raw data for LC-MS analysis of GAGs are available at GlycoPOST [84] under project ID GPST000731. Raw sequencing reads and results of the RNA sequencing and CRISPR screening experiments will be made available upon publication. Any data supporting the analyses in the manuscript are available from the corresponding author upon reasonable request.

## Supplementary Information

Supplementary tables and figures are provided in the Supporting Information.

## Acknowledgements

We thank IBEX Technologies for their in-kind donation of heparin lyase enzymes. We thank Dr. Jeffrey Esko for insightful discussions supporting this project. RNA sequencing and CRISPR amplicon sequencing were conducted at the IGM Genomics Center, University of California, San Diego, La Jolla, CA. This publication includes data generated at the UC San Diego IGM Genomics Center utilizing an Illumina NovaSeq 6000 that was purchased with funding from a National Institutes of Health SIG grant (#S10 OD026929). Qubit quantification and MiSeq analysis of Calabrese sgRNA plasmid library were performed at the Georgia Genomics and Bioinformatics Core (GGBC, UG Athens, GA, RRID:SCR_010994). We thank Juan Bustamente and Julie Nelson at the UGA CTEGD Cytometry Shared Resource Laboratory for their help with FACS sorting experiments and additional advice. R.J.W. is supported by grants from NIH (R35GM150736 and R21HL167091). K.M. is supported by grants from NIH (R01GM154846). Z.W. and J.L. are supported by grants from NIH (R44GM142304 and R01AG087305). The carbohydrate analyses performed at the CCRC were supported by NIH grant R24GM137782 to P.A. Certain figure panels (Figs. 3A, 6A) were created in BioRender.

## Author Contributions

Unless otherwise noted, J.C.M. performed the experimental work and analyzed the data. H.T. and C.N. characterized activation cell lines and performed immunoblotting, flow cytometry, and qPCR experiments. C.H., D.C., and K.M. expressed, purified, and characterized recombinant proteins. A.B. generated, processed, and analyzed the LC-MS HS compositional data. Z.W. and J.L. generated, processed, and analyzed the LC-MS/MS HS tetrasaccharide and CS/DS compositional data. J.C.M. and R.J.W. wrote the paper, with input from all listed co-authors.

