## Supplementary Information for "CRISPR activation screens identify core protein-dependent regulation of heparan sulfate sulfation and ligand specificity"

**This PDF file includes:**

Supplementary Tables 1-6
Supplementary Figures 1-4

**Supplementary Table 1:**
HS disaccharide composition for HS chains isolated from recombinant eSDC1 and eSDC2

| **HS Disaccharides** | | **Abundance (% Total Disaccharide)** | |
|---|---|---|---|
| **Structure Code[a]** | **Unit Formula[b]** | **eSDC1 (% Total HS)** | **eSDC2 (% Total HS)** |
| D0A0 | ΔUA-GlcNAc | 69.20 ± 1.04 | 71.44 ± 0.60 |
| D2A0 | ΔUA2S-GlcNAc | 1.13 ± 0.03 | 0.82 ± 0.02 |
| D0A6 | ΔUA-GlcNAc6S | 2.40 ± 0.09 | 1.19 ± 0.14 |
| D2A6 | ΔUA2S-GlcNAc6S | 0.10 ± 0.02 | 0.05 ± 0.00 |
| D0S0 | ΔUA-GlcNS | 14.73 ± 0.68 | 14.71 ± 0.26 |
| D2S0 | ΔUA2S-GlcNS | 9.59 ± 0.61 | 10.13 ± 0.46 |
| D0S6 | ΔUA-GlcNS6S | 2.01 ± 0.06 | 1.20 ± 0.11 |
| D2S6 | ΔUA2S-GlcNS6S | 0.84 ± 0.21 | 0.46 ± 0.10 |

[a] The disaccharide structure code is described in (Lawrence, et al. *Nat. Methods* 2008)
[b] ΔUA = 4,5-unsaturated uronic acid

**Supplementary Table 2:**
Distribution of HS disaccharides for HS chains isolated from recombinant ectodomain SDC1 (eSDC1) and SDC2 (eSDC2)

| **HS Sulfates/disaccharide** | **Abundance (% Total Disaccharide)** | |
|---|---|---|
| | **eSDC1** | **eSDC2** |
| N-Ac | 73.71 ± 1.14 | 73.88 ± 0.46 |
| N-$SO_3$ | 25.15 ± 1.11 | 25.30 ± 0.46 |
| 2-*O* $SO_3$ | 13.67 ± 0.50 | 12.66 ± 0.47 |
| 6-*O* $SO_3$ | 5.35 ± 0.35 | 2.90 ± 0.15 |

**Supplementary Table 3:**
HS tetrasaccharide composition for HS chains isolated from recombinant eSDC1 and eSDC2

| **HS tetrasaccharides** | | **Abundance (pg / mg protein)** | |
|---|---|---|---|
| **Structure Code[a]** | **Unit Formula[b]** | **eSDC1** | **eSDC2** |
| D0N6-G0S9 | ΔUA-GlcNAc6s-GlcA-GlcNS3S6S | 4.75 ± 0.50 | 1.25 ± 0.50 |
| D0S6-G0S9 | ΔUA-GlcNS6S-GlcA-GlcNs3S6S | 18.25 ± 4.50 | 7.25 ± 5.25 |
| D0S6-I2S9 | ΔUA-GlcNS6S-IdoA2S-GlcNS3S6S | 9.25 ± 2.87 | 8.5 ± 1.29 |
| D0S0-I2S3 | ΔUA-GlcNS-IdoA2S-GlcNS3S | 79.50 ± 5.80 | 109.50 ± 6.45 |
| D2S0-I2S3 | ΔUA2S-GlcNS-IdoA2S-GlcNS3S | 53.50 ± 14.55 | 78.50 ± 9.85 |

[a] The tetrasaccharide structure code is described in (Lawrence, et al. *Nat. Methods* 2008)
[b] ΔUA = 4,5-unsaturated uronic acid

**Supplementary Table 4:**
CS/DS disaccharide composition for CS/DS isolated from recombinant eSDC1 and eSDC2

| CS/DS Disaccharides | | Abundance (% Total Disaccharide)[c] | |
|---|---|---|---|
| Structure Code[a] | Unit Formula[b] | eSDC1 (% Total CS) | eSDC2 (% Total CS) |
| D0a0 | ΔUA-GalNAc | 11.12 ± 0.04 | 18.77 ± 0.09 |
| D0a4 | ΔUA2S-$GalNH_2$ | 51.61 ± 0.27 | 49.26 ± 0.23 |
| D0a6 | ΔUA-GalNS | 36.78 ± 0.27 | 31.66 ± 0.31 |
| D2a10 | ΔUA-GalNAc6S | 0.50 ± 0.00 | 0.31 ± 0.00 |

[a] The disaccharide structure code is described in (Lawrence, et al. *Nat. Methods* 2008)
[b] ΔUA = 4,5-unsaturated uronic acid

**Supplementary Table 5:**
sgRNA sequences used in this study

| Gene Target | Forward guide sequence (5'-3') | Reverse guide sequence (3'-5') |
|---|---|---|
| *HS3ST1* | AGGGAGAGCGCGTTGGGCAG | CTGCCCAACGCGCTCTCCCT |
| *SDC1* | GGCGTTCCGAAGGGGCCGGG | CCCGGCCCCTTCGGAACGCC |
| *SDC2* | AGAAGCAGGCTCAGGAGGGA | TCCCTCCTGAGCCTGCTTCT |
| *SDC3* | GCGGCGGCGCACGCTTCCTG | CAGGAAGCGTGCGCCGCCGC |
| *SDC4* | CCCCGCCCGGAATTCCCCAG | CTGGGGAATTCCGGGCGGGG |

**Supplementary Table 6:**
Primer sequences used for quantitative PCR experiments

| Gene | Forward primer sequence (5'-3') | Reverse primer sequence (5'-3') |
|---|---|---|
| *YWHAZ* | CCTGCATGAAGTCTGTAACTGAG | GACCTACGGGCTCCTACAACA |
| *SDC1* | ACGAAGGCAGCTACTCCTTG | GTTTGGTGGGCTTCTGGTAG |
| *SDC2* | GCTGTTGGTGTATCGCATGA | ACTGGATGGTTTGCGTTCTC |
| *SDC3* | GCCACCACTGCTGTTATAAGG | ACTGTGGTCAGTGGGAGAGG |
| *SDC4* | ACTGTGGTCAGTGGGAGAGG | GCTGCCTTCATCCTTCTTCTT |
| *HS3ST1* | AACGAGGTCCACTTCTTCGA | GCATCTGGCTGAGGTACCAG |
| *HS3ST3A1* | CGGAAGTTCTTGCTGATGCT | CTCGGCCAGGCAGTAGAA |
| *HS3ST3B1* | CCGGTGAGGAGGAAGCTC | GGCGCACGAGTACAGGAA |
| *HS3ST5* | CAATTTCATGTCGTCGATGG | AGGAACTTCTCCACGAGCTG |
| *NDST1* | TCCTGCTGTTCATCTTCTGC | CTCGCTTCCAGCCATATAGG |
| *HS6ST2* | TACACTGGCGATGACTGGTC | GCGGTTGTTGGCTAGATTGT |

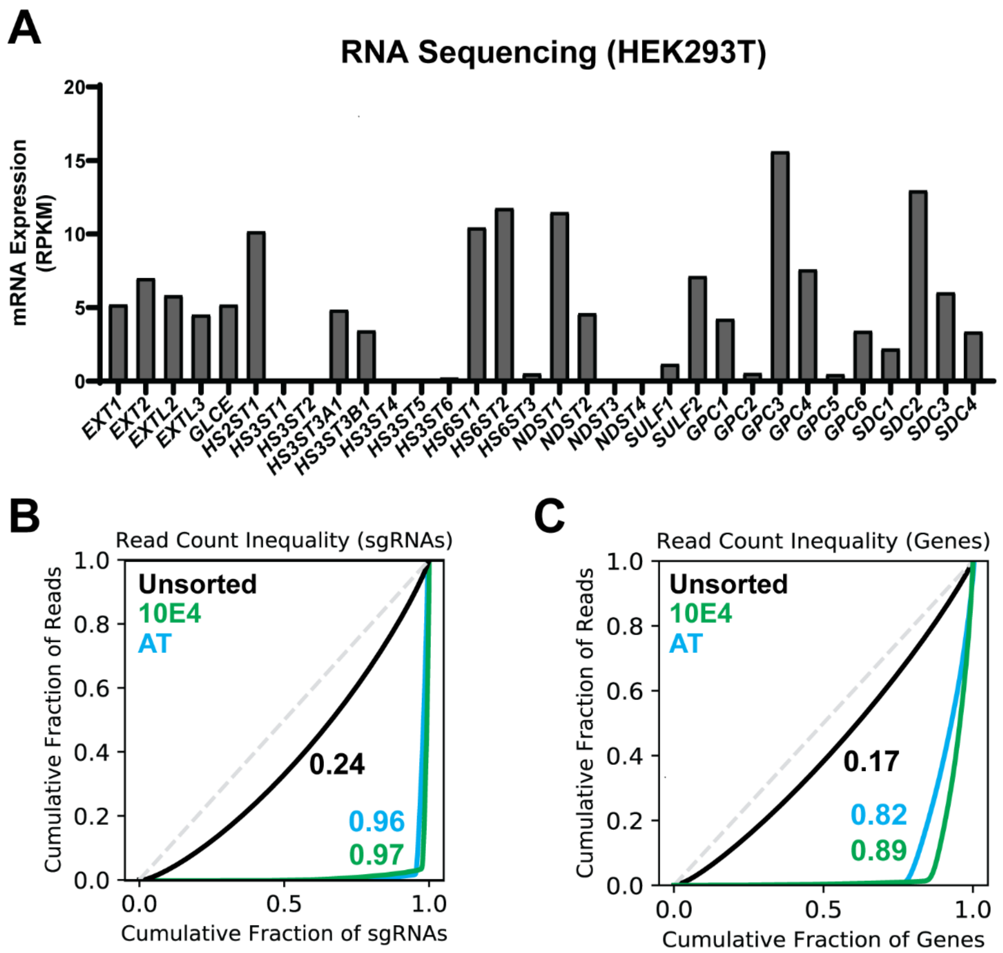


**Supplementary Figure 1. HEK293T RNA sequencing data for HSPG genes and CRISPRa screening read distributions. (A)** RNA Sequencing of HEK293T WT cells. RPKM values shown represent an average of biological duplicates (n = 2 independent biological replicates). Lorenz curves showing the distribution of **(B)** sgRNA and **(C)** gene sequencing reads in unsorted control cells versus 10E4-, and AT-sorted cell populations. Numbers represent Gini coefficients (0: reads cover sgRNA/gene library evenly, 1: reads cover only a single sgRNA/gene).

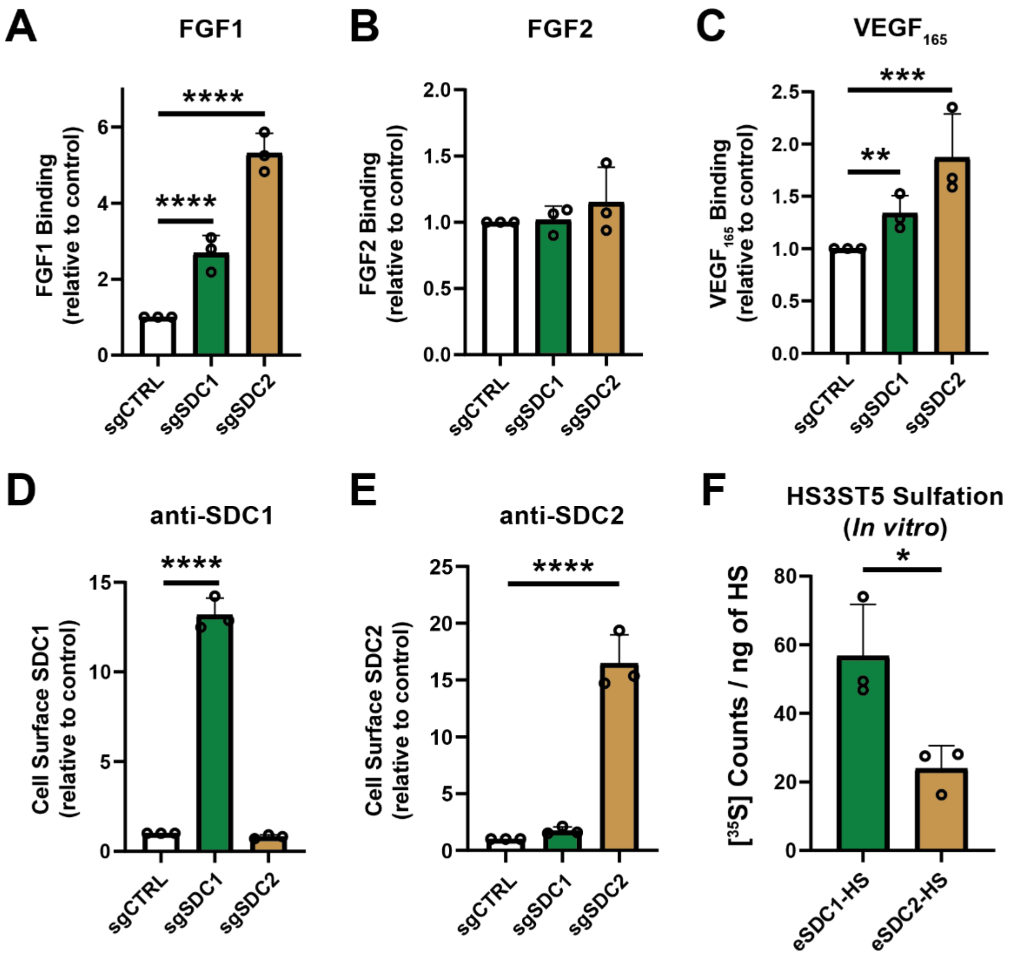


**Supplemental Figure 2. Characterization of SDC1 and SDC2 CRISPRa cell lines.** Flow cytometry analyses of **(A)** FGF1, **(B)** FGF2, and **(C)** $VEGF_{165}$ cell surface binding for SDC1 and SDC2 activation lines compared to sgCTRL cells. **(D-E)** Flow cytometry analyses of cell surface SDC1 and SDC2 levels in activation lines versus control cells. **(F)** [$^{35}$S] counts normalized to HS mass input for in vitro HS3ST5 sulfation reactions. Data are presented as mean ± SD (n = 3 independent experiments), ****$p$<0.0001, ***$p$<0.001, **$p$<0.01, *$p$<0.05 by two-sided t-test.

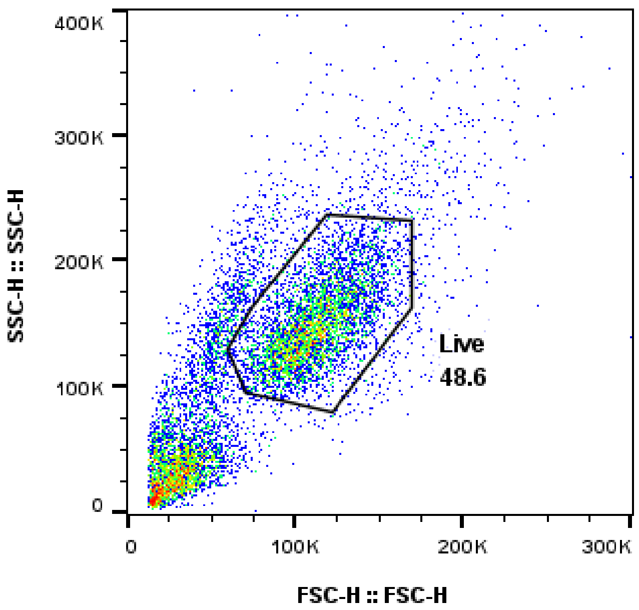


**Supplementary Figure 3. General flow cytometry gating strategy.** Cells were gated based on forward and side scattering for analysis of flow cytometry data.

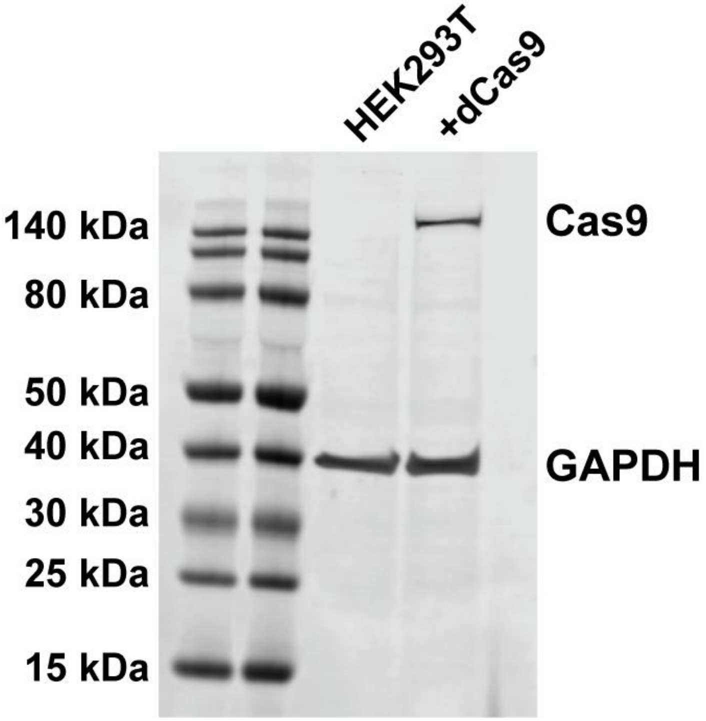


**Supplementary Figure 4. Source Data.** Uncropped western blot source image for Figure 1B.
